# Activity patterns during the mating season predict sex-biased infections in an emerging fungal disease

**DOI:** 10.1101/2024.09.02.609945

**Authors:** Macy J. Kailing, Joseph R. Hoyt, J. Paul White, Jennifer A. Redell, Heather M. Kaarakka, Kate E. Langwig

## Abstract

1. Mating dynamics can govern species impacts from rapid global change by influencing population rates of growth and adaptation, as well as individual traits that affect mortality risks from novel pressures.
2. Here, we examined sex differences in activity of *Myotis lucifugus* during their mating season, which coincides with exposure to the lethal fungal pathogen (*Pseudogymnoascus destructans*) that causes white-nose syndrome. We expected differences in activity between sexes to modify seasonal *P. destructans* dynamics as the pathogen can replicate only at the cool temperatures at which bats hibernate.
3. We used passive antenna systems installed at the entrances of hibernacula and PIT tags to characterize activity patterns of bats. We also measured pathogen loads on bats during the autumn mating and early hibernation periods to assess how infection changed according to host phenology.
4. We found that females spent fewer days active during autumn, arrived after males, and were primarily active on the warmest nights. Males remained highly active throughout the mating season and later in autumn than females. Importantly, differences in phenology corresponded to higher pathogen loads on females during early hibernation as male activity, and thus warm body temperatures, inhibit pathogen growth.
5. Differences in activity between sexes and in the transition from swarm to hibernation likely reflects males maximizing their mating opportunities while females conserve energy to meet the cost of spring migration and reproduction. More broadly, our results show how activity during the mating season and phenology can contribute to sex-biased impacts of a novel disease and highlight the value of understanding species’ mating systems to anticipate the impacts of environmental change.

## 1. Introduction

Global change has imposed novel pressures that threaten many species with extinction, including habitat loss, pollution, overutilization and emerging infectious disease (Dirzo et al., 2014; Mazor et al., 2018; Otto, 2018; Tilman et al., 2017). The resilience of populations to global changes can vary based on the species’ life history, including the mating system (Capdevila et al., 2022; Péron, 2013; Plesnar-Bielak et al., 2012; Sæther et al., 2004). Mating systems can govern how quickly populations recover from and adapt to novel stressors by influencing demographic and evolutionary processes, such as rates of recruitment and genetic mixing (Bessa-Gomes, Legendre, & Clobert, 2004; Legendre et al., 1999). Further, mating strategies, characterized by the collection of activities and behaviors used by individuals to monopolize and secure mates, can vary within species and be a substantial source of trait variation (Bradbury & Vehrencamp, 1977; Clutton-Brock, 1989; Shuster & Wade 2003). Accordingly, traits expressed during mating periods are often shared within demographic classes (e.g., sex, age, social status, etc.) and can result in selective mortality that can restructure or destabilize populations (Clutton-Brock et al., 2002; Grüebler et al., 2008; Martin et al., 2022; Oro, Álvarez, & Velando, 2018; Rosa et al., 2019; Wells et al., 2017). Therefore, understanding how aspects of the mating system influence interactions between individuals and novel pressures should improve the ability to predict responses of populations to rapid changes.

Sexes use different strategies for maximizing their fitness that give rise to variation in morphology, behaviors, or physiology (Kolata, 1977; Promislow, Montgomerie, & Martin, 1992; Trivers, 1972), which collectively can affect their mortality risk (Promislow, 1997). In many species where sexual selection leads to dimorphic traits between sexes, males invest in quantity of offspring through copious mating, whereas females, who are more limited by gametic production, invest in ensuring survival of few high-quality offspring (Bateman, 1948; Lehtonen, Parker, & Schärer, 2016). Accordingly, males may be more likely to engage in risky behaviors to increase their competitive mating advantages whereas females, whose fitness relies on surviving to carry and care for young, may be more risk-averse (Bateman, 1948; Bonduriansky et al., 2008; Promislow, 1997). Additionally, asynchrony between sexes in their reproductive investments can also contribute to differences in how females and males respond to the same environmental pressures (Cornwallis & Uller, 2010; Forsgren et al., 2004; Kasumovic et al., 2008), including when their peak vulnerability to stressors occurs (Williams et al., 2022). While sex specific responses to novel pressures are expected, the proximate factors that influence differences in impacts between females and males are overlooked for many novel threats (Connallon, Débarre, & Li, 2018; Connallon & Hall, 2018).

Novel pressures like infectious diseases frequently interact with mating systems resulting in differential impacts to sexes (Herrera & Nunn, 2019; Stephenson, 2019; Stephenson et al., 2016; Zuk, 2009; Zuk & McKean, 1996) that can influence the severity of epidemics and coevolution between hosts and pathogens (Hall & Mideo, 2018). Typically, male behaviors or physiological states optimize fitness at the costs of higher pathogen exposure or growth, including territory defense or aggression, dispersal, and reduced immunity (Ferrari et al., 2003; Grear, Luong, & Hudson, 2012; Luong et al., 2010; Sauer et al., 2024; Stephenson, 2019). For example, adult male badgers had higher contacts with conspecifics and higher risks of acquiring tuberculosis compared to females (Delahay et al., 2006; Graham et al., 2013; McDonald et al., 2014). Male-biased disease is more frequently documented, but female-biased disease has been shown to arise seasonally (Cattadori et al., 2005; Sweeny et al., 2021). For example, during parturition adult female European rabbits had the sharpest peak in nematode intensities relative to non-reproductive females and males (Cattadori et al., 2005), illustrating that the magnitude and direction of sex-biases can vary temporally to affect the dynamics between hosts and pathogens (Cattadori et al., 2005; Sweeny et al., 2021). However, the behavioral and physiological mechanisms contributing to sex-biases in impacts from introduced pathogens are often unclear, thus limiting opportunities for management.

White-nose syndrome (WNS) is a recently introduced disease of bats that is caused by the psychrophilic fungal pathogen, *Pseudogymnoascus destructans*, and has led to population declines exceeding 95% (Frick et al., 2015; Langwig et al., 2012; Langwig et al., 2016). Euthermic bats that are active on the landscape maintain higher body temperatures (37-41°C) (O’Shea et al., 2014) that are above the upper critical limit of *P. destructans* growth (20°C) (Verant et al., 2012). However, when bats reduce their body temperature during torpor (short periods of inactivity that comprise hibernation) to near ambient (1-16°C) (Perry, 2013; Webb, Speakman, & Racey, 1996) the pathogen is able to grow, making it dependent on seasonal hibernation cycles (Jackson, Willcox, & Bernard, 2022; Jackson et al., 2022; Langwig et al., 2015; Langwig et al., 2021). Pathogen exposure occurs during autumn when bats return annually to contaminated hibernation sites (Hoyt et al., 2020; Laggan et al., 2023) but infection and mortality peak over winter (Hoyt, Kilpatrick, & Langwig, 2021; Langwig et al., 2015; Reeder et al., 2012).

The mating system for temperate bats impacted by WNS is polygynandrous and described as a swarm (Fenton 1969, Bradbury & Vehrencamp 1977). Swarming occurs during autumn when large numbers of individuals aggregate near hibernacula entrances, then enter the site to form smaller groups (typically pairs or triads), and copulate after landing on walls or in crevices (Fenton, 1969; Fraser & McGuire, 2023; Thomas, Brock Fenton, & Barclay, 1979; Wai-Ping & Fenton, 1988). While females mate during this period, they store sperm through hibernation until spring emergence, when they migrate to summer maternity colonies, become pregnant, and rear pups cooperatively with other females (Fenton & Robert, 1980; Kurta & Kunz, 1987; Wimsatt, 1942, 1944, 1945). Males segregate from females during parturition and pup rearing (Davis & Hitchcock, 1965; Fenton & Robert, 1980). To that end, females’ reproductive investments are primarily during spring and summer, and are more extensive in magnitude and duration than males, whose reproductive investments are highest during autumn (Racey & Entwistle, 2000; Fig 1). While both sexes mate during autumn, asynchronous reproductive investments between females and males could lead to extensive behavioral and physiological traits that contribute to disease impacts. Specifically, the congregation of sexes at contaminated hibernacula during autumn and hibernation provide a unique opportunity to explore how sex differences in activity during the mating and pathogen exposure period influences infection intensity between sexes.

**Figure 1.**
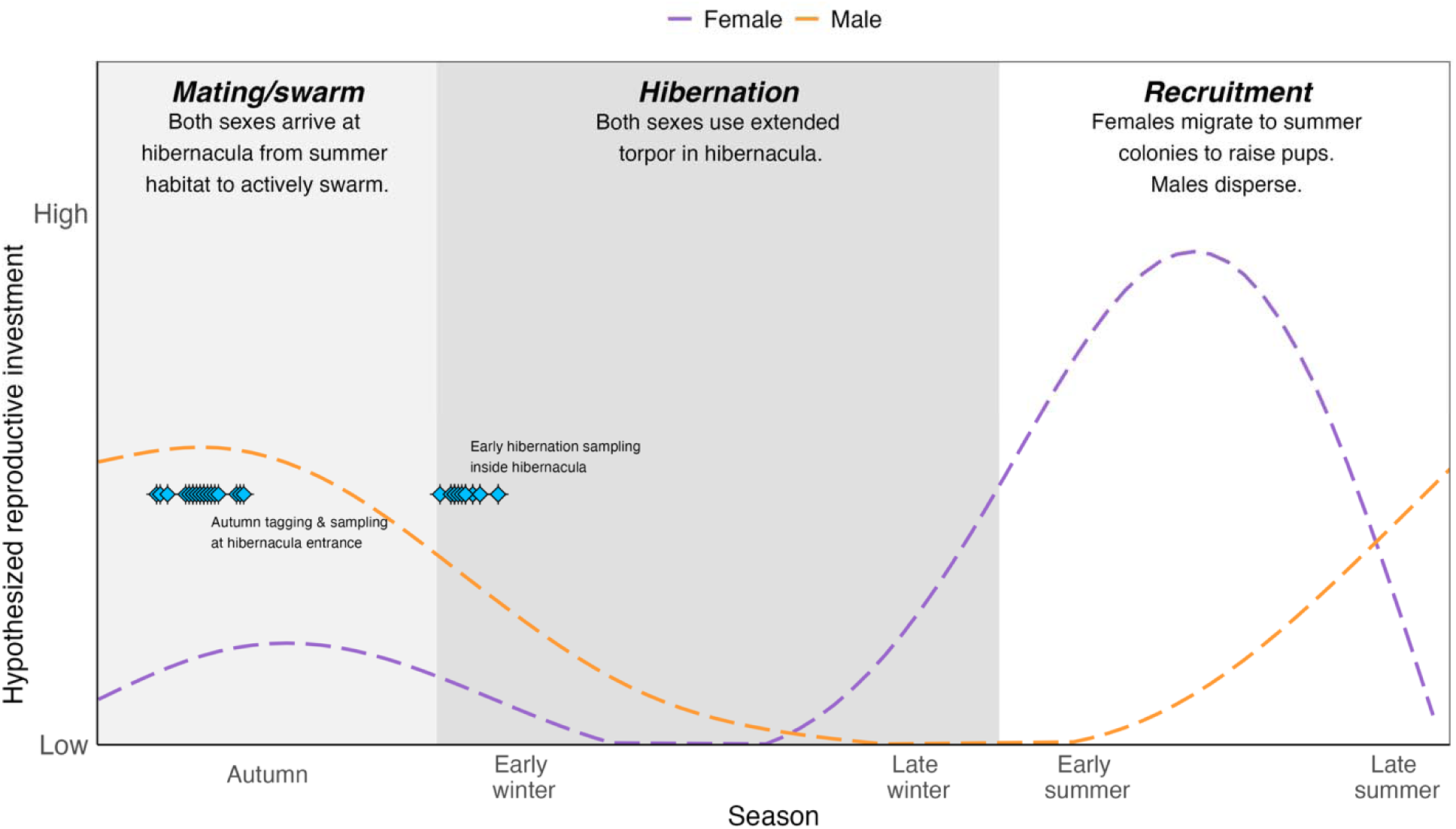
Hypothesized reproductive investment of females and males with different reproductive strategies developed from Fenton & Robert, 1980, Racey & Entwistle, 2000, and Wimsatt, 1945. Males are expected to invest most towards their reproductive fitness during autumn when they mate with many females. Females likely require less energy during the autumn swarm, but considerable investment is made following hibernation as they singly parent their young in maternity colonies and require energy for gestation, parturition, and lactation. Light blue points correspond to the tagging and sampling dates during the seasons of autumn mating and early hibernation as described in Methods.

Temperate bats reduce activity seasonally and use torpor to conserve energy even outside of winter hibernation (Alston et al., 2022). However, females and males balance torpor and activity differently throughout the year (Grinevitch, Holroyd, & Barclay, 1995; Hoff et al., 2023). These sex specific differences in activity and torpor could strongly influence WNS dynamics if sex-specific torpor use influences growth of the cool-temperature dependent fungus. Yet, empirical investigations are limited by the challenge of obtaining individually based data on activity patterns through time. Previous research has shown that females can be more infected and suffer greater impacts (Grieneisen et al., 2015; Kailing et al., 2023), but see Johnson et al., 2014. However, the mechanisms driving the difference between sexes remain unknown. We hypothesize that activity during the autumn mating season drives highly disparate infections in early hibernation since the pathogen is unable to grow on bats when they are active. As such, we investigated how traits associated with mating strategy, evaluated here based on activity during the mating season, vary between females and males and contribute to sex-biased infection intensity during seasons important for reproduction (autumn mating) and disease (autumn mating and early hibernation). Specifically, we aimed to characterize the active period at swarm locations (i.e., hibernacula), examine environmental and host factors associated with activity during the mating season, and explore links between phenology and infection intensity.

## 2. Methods

### 2.1 Field monitoring

We studied the activity of *Myotis lucifugus* in Wisconsin, USA during the autumn mating season each year from 2020 to 2023 at three large hibernacula within Pierce and Dodge counties, consisting of 10 total entrances. We considered July 01 to November 01 as the autumn mating period as activity patterns at the hibernacula are likely to be motivated by breeding. We used this relatively broad timeframe to avoid missing the earliest arriving or latest swarming bats. At every entrance, we installed an RFID system consisting of an antenna with an automated data logger (IS10001; Biomark) that recorded a time-stamped datapoint each time a passive integrated transponder (PIT) tag was in proximity. Each system was scheduled to run continuously by a direct or solar source of power. Systems were programmed to record unique tags every minute to help to maximize detections as >1 tag cannot be simultaneously recorded and bats sitting on top of antenna would otherwise block detections of all other individuals. In addition, systems recorded test tags at least once every 12 hours enabling us to determine when systems were operational.

### 2.2 PIT tagging and measuring pathogen quantity

We captured bats at each site during autumn (between August 15 and September 02, Fig. 1) using a harp trap (Harptrap; Faunatech Austbat) and mist-nets (Polyester 38mm mesh; Avinet) near the entrances where individuals were active. For every individual, we weighed, injected a PIT tag, categorized sex based on external morphology, and attached an aluminum-lipped band (2.9 mm; Porzana, Icklesham, UK) as secondary identification. Capture efforts at each site concluded after one night, unless the target number of bats (approximately 100) was not achieved, in which we trapped a second night.

We also sampled bats in early hibernation (between November 01 and November 17, Fig. 1) at the same sites by removing them from their roosting locations on the walls and ceilings of the hibernacula. We walked sections of the hibernacula to collect a stratified sample of up to 20 bats. As in autumn, we recorded sex based on external morphology of every individual and attached a band. We collected an epidermal swab from all the bats that were sampled in autumn or early hibernation with a standardized swabbing technique that is used widely to assess pathogen presence across seasons (Frick et al., 2015; Langwig et al., 2015). Immediately upon collection, we stored swabs in RNAlater. We tested swab samples for *P. destructans* using Qiagen DNeasy Blood and Tissue Kits (Qiagen, USA) to extract fungal DNA followed by qPCR to quantify fungal loads (Langwig et al., 2015; Muller et al., 2013; McGuire et al., 2016). All bat handling procedures were granted ethical approval by the Institutional Animal Care and Use Committee at Virginia Tech (IACUC Protocols #20-150, #23-197), field work was conducted under permits granted by the Wisconsin Department of Natural Resources (Endangered and Threatened Species Permits #886, #882), and decontamination procedures followed United States Fish and Wildlife recommendations.

### 2.3 Characterizing individual activity and climate variables

We determined individuals’ active periods (i.e., the timing in which mating could occur based on a bat being active at the swarm location) from PIT tag detections. An individual was considered active if it was detected at any entrance to the site at least once in a night because bats commonly visit multiple entrances throughout the night. We classified bats that were detected between the hours of 00:00 and 07:00 as active on the previous date because bats are nocturnal, and thus, their active date begins and ends at sunset and sunrise, respectively. We only included bats detected on at least two nights per year to properly categorize the detections as being the bats’ first or last active date within the same year to capture the apparent swarm period. For each bat observed within a year, we assigned a start swarm date as the first date detected after July 01 and an end swarm date as the last date detected before Nov 01. Start dates were only assigned for individuals that returned from a previous year as we could not confirm when bats first arrived at hibernacula if they were untagged. We were able to assign an end swarm date to all bats, including those tagged in the same year, because the last detection always followed the date they were captured and tagged. *Myotis lucifugus* can only be reliably aged as juveniles (young of the year) or adults and all bats that return from a previous year are considered adults. Therefore, we only included adults in all analyses.

To assess environmental factors associated with activity during the mating period, we extracted climate data using the R package ‘daymetr’ (Hufkens et al., 2018) and GPS coordinates of hibernation sites, which provided continuous, grid-based estimates of daily weather variables based on location. Because most bats begin nightly activity at sunset before temperatures reach an overnight low, the warm temperatures prior to sunset are relevant for activity. Therefore, we calculated a mid-point temperature (Jackson et al., 2022) for each date as: (minimum temperature on the following day + maximum temperature of current day)/2 to incorporate the warm and cool temperatures that could influence bat activity and because mean temperatures were not available in the climate data.

### 2.4 Statistical analyses

#### 2.4.1 Summary

We tested our hypotheses using generalized (i.e., GLMM) and linear (i.e., LMM) mixed effects models to characterize activity throughout the autumn mating period and pathogen quantity by sex with R packages ‘lme4’ (Bates et al., 2015) and ‘glmmTMB’ (Brooks et al., 2017). For each model, we tested variables based on *a priori* hypotheses and evaluated their significance using ANOVA tables for statistical model objects from the ‘car’ package (Fox & Weisberg, 2019). We combined site and year of sampling (site-year) to use as a random effect in all models and randomize the intercepts of annual sampling events. Condensing to site-year was more relevant for our analyses than using site and year separately because our sites are relatively far apart and years vary in time since pathogen arrival. In analyses where individuals were observed repeatedly in the same site-year, we included band or PIT tag ID as an additional random effect. We compared candidate and null models using Akaike Information Criterion (AIC) to evaluate relative quality (difference in AIC scores |> 2|; See Appendix 4.6.2). We also visually checked model assumptions, such as the residual distribution of linear models, and conducted k-fold cross-validation of logistic models to check predictive accuracy (Appendix 4.6.3) using the ‘performance’ package (Lüdecke et al., 2021). We also used the ‘performance’ package (Lüdecke et al., 2021) and model outputs to visualize normality and verify the structure of our random effects. Because many more males were detected than females, we also generated a balanced dataset between sexes to check that the results informed by our male-biased dataset were consistent and not due to the sampling bias. We truncated our full dataset by randomly selecting the number of male observations equivalent to the number of female observations from all the male observations (see Supplemental Methods). We extracted model estimates using R package ‘emmeans’ (Lenth, 2022) to report final models. See Supplemental Table 1 for overview of all analyses as well as Appendix with code for all statistical models, output, and model validation.

#### 2.4.2 Dates of autumn mating activity

We determined individually based active periods at swarm locations using LMMs. First, we estimated when individuals of each sex started activity during autumn using the first date an individual was detected at the hibernacula as a continuous response variable, with sex as a fixed effect (F|M), and site-year and individual as random effects. Then, we used the same model structure, but with last date detected as a response to estimate when individuals of each sex ended activity. Prior to analyses, we converted dates to continuous monthly units with decimal days by dividing the day of each month by 31 and adding it to numeric month. We then scaled the dates to start at 0 on July 1. For example, July 15 corresponded to 0.5.

#### 2.4.3 Effects of temperature and body condition on nightly activity

To test whether environmental conditions differently affected nightly activity between sexes, we used a GLMM with a binomial distribution to test the probability of an individual being active (=1) or undetected (=0) with temperature and sex as interactive fixed effects and individual and site-year as random effects. For the response variable, we treated each bat as a series of binomial trials using all the possible dates between the first and last detection where an individual was detected (=1) or undetected (=0). We excluded nights where the minimum temperature was less than 10°C to minimize outlier effects as bat activity is greatly reduced (Gorman et al., 2021).

We also tested the effect of body condition on nightly activity to assess whether energetics might contribute to differences in activity between males and females. As we could only associate mass to bats at capture, for this dataset, we used the activity of bats in their first year tagged. We used a GLMM with a binomial distribution to test the probability of a bat being active. Like the above analysis, our response variable was detection probability, with each bat treated as a series of binomial trials where it was either detected or undetected. We used sex, mass, and temperature as interactive fixed effects and included site-year and individual as a random effect.

#### 2.4.4 Effects of length of active period and temperature on last detection

We examined how the number of days since arriving to the hibernaculum and temperature affected whether sexes transition from being active to hibernating to understand how host factors (time spent active) or environmental conditions (temperature) influence phenology. First, we used a GLMM with a binomial distribution that predicted whether individuals stopped activity for the year. For the response variable, detections of each bat were treated as a series of binomial trials where an individual was detected for the last time (=1) or was detected on a subsequent date (=0). We included interactive effects of sex and time since their first detection (to standardize differences in start date) with site-year and individual as random effects. For days since first detection, we binned dates to 3-day intervals and counted the number of intervals that passed since their arrival to the hibernacula. Accordingly, we only included individuals that we were able to assign start dates (i.e., returned from a previous year). Date intervals were used to overcome sparsity in the number of females that are active each night.

To also examine the effect of temperature on the conclusion of autumn activity, we used a GLMM with a binomial distribution to test a similar response as above where each bats’ detections were treated as a series of binomial trials and an individual was detected for the last night (=1) or was detected on a subsequent night (=0). We included sex and temperature as interacting fixed effects and random effects of site-year and individual. We tested length of swarm period and temperature interacting with sex in separate models given collinearity between the two predictors.

#### 2.4.5 Seasonality and host phenology effects on pathogen quantity

To understand infection dynamics according to host phenology, we tested differences in pathogen quantity between sexes and across seasons using a LMM with log_10_ *P. destructans* loads as our response variable, with sex and date (in monthly units as above) as fixed effects, and site-year as a random effect. Because we did not resample the same bats between autumn and early hibernation within the site-year, we did not include individual ID as a random effect. We explored associations between phenology and infection intensity further by testing the effect of the active period on early hibernation pathogen quantity. We used a LMM to test log_10_ *P. destructans* loads as a response variable with median last date of activity as a fixed effect. We included site-year as a random effect. We calculated the median date of last detection in autumn for both sexes for each site and year (i.e., the end of swarm activity and assumed start of hibernation) and assigned it to individuals that we sampled for pathogen quantity in early hibernation of the respective site and year. We used the median date of last detection to minimize the weight of outlier individuals that may leave the hibernacula late in autumn but are not representative of swarm activity. We also performed this analysis at the population level using the mean early hibernation pathogen quantity from each site and year as the response. We used these approaches because genetic analyses indicate that swarming and hibernating populations at a site are generally comprised of the same individuals (van Schaik et al., 2015) but relocating and sampling the same individuals across seasons is rarely achieved, in part due to the complexity of passages and high ceilings of hibernacula. We also evaluated autumn temperature as a predictor of pathogen quantity.

## 3. Results

We characterized autumn activity of 148 (by site: 37, 40, 71) female and 608 (by site: 225, 155, 228) male *M. lucifugus* and measured pathogen quantity during capture on 168 (by site: 56, 63, 49) female and 416 (by site: 151, 130, 135) males between 2020 and 2023. See Supplemental Table 1 for overview of all analyses and Appendix 4.7 for full sample lists used in each analysis by site, years, and sex. We also provide a visual representation of the full dataset in Supplemental Fig. 1 (Appendix 4.1). All statistical results are reported with males as the intercept or reference level and thus coefficient ± SE are in reference to the value above or below the male references. Females started swarm activity later (Female coef ± SE = 0.249 ± 0.08, t = 3.25, p-value = 0.001; Fig. 2, top panel; Appendix 1.1) but also ended activity earlier than males (−0.387 ± 0.06, t = −6.29, p-value < 0.001; Fig. 2, top panel; Appendix 1.2). Male activity fully encompassed the active period of females (Fig. 2, bottom panel; Appendix 1.3) and the number of individuals detected was overwhelmingly male biased throughout autumn, given their higher abundance and activity overall (Supplemental Fig. 1; Appendix 4.1), and was consistent between sampling events (Supplemental Table 2; Appendix 4.6.1).

**Figure 2.**
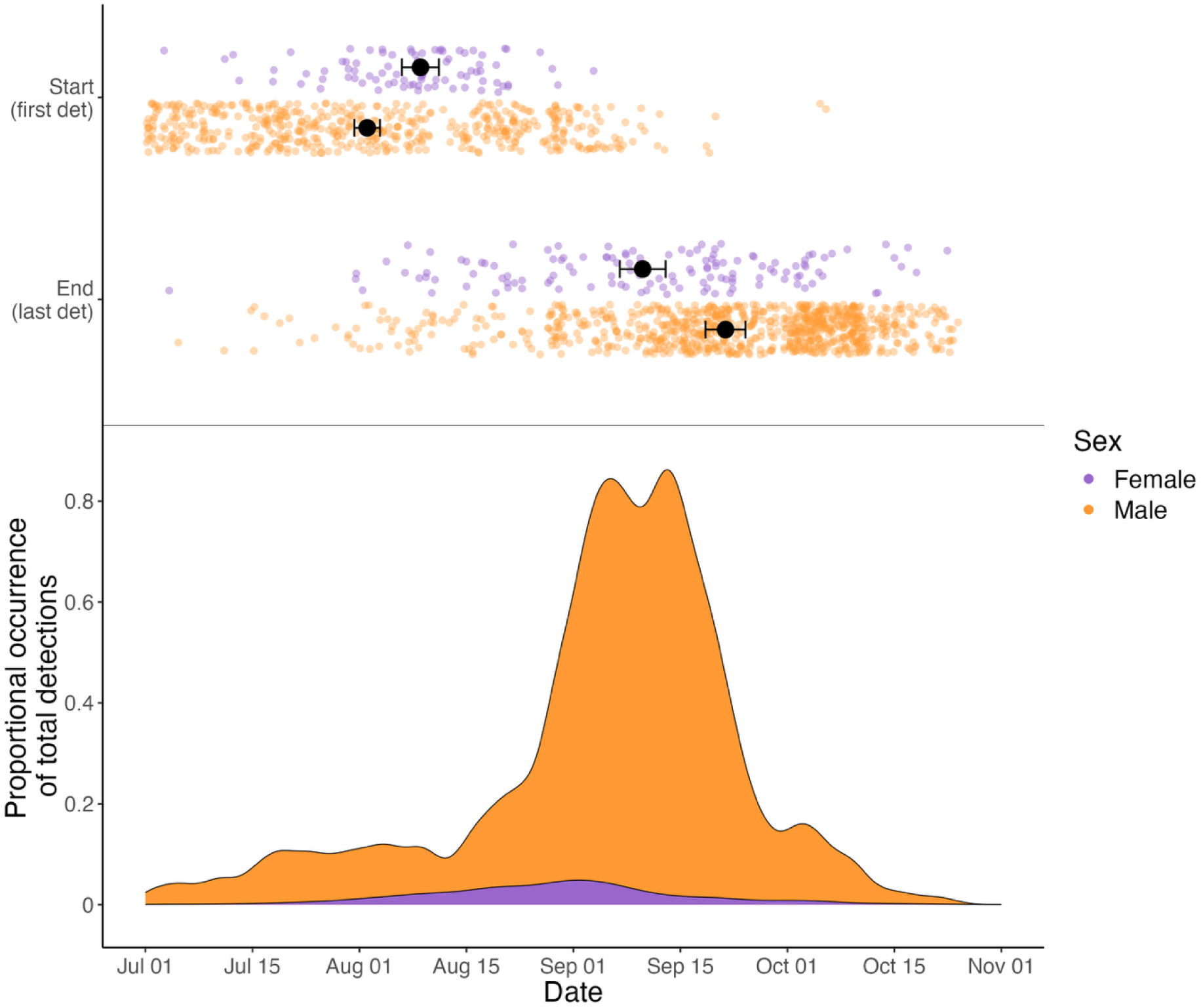
**Top:** Differences in the autumn active period by sex. Each point represents the date of starting (first detection) and ending (last detection) activity during autumn mating for an individual bat. Black circles represent model estimated dates with standard errors shown with horizontal bars. **Bottom:** Distribution of activity by sex. Each density plot shows the proportion of all detections grouped by sex that occurred on each date. An individual bat can be represented on multiple nights if activity occurred. Asymmetry in the peak of male activity (based on the total number of observations) relative to their estimated active period in the top panel illustrates visually that while most males arrive early in autumn, they maintain activity into later swarm dates.

Within their respective active periods, the probability of being active in a night or not for each sex was differently affected by temperature (detected=0|1; Female*temperature = 0.151 ± 0.03, t = 4.97, p-value < 0.001, Fig. 3A; Appendix 2.1), such that the probability of females being active was lower overall but increased more quickly with increasing temperatures (Temperature slope estimated using ‘emtrends’: Female slope = 0.152 ± 0.03 vs Male slope 0.001 ± 0.01, Fig. 3A; Appendix 2.1), notably above 16°C. An interaction between sex and temperature was also evident when using a balanced dataset of males and females (see Supplemental Statistical Methods, Supplemental Fig. 2A; Appendix 4.2.1). We found that the effect of body condition on activity differed between males and females, with light females less active at cooler temperatures and more active at warmer temperatures, whereas all males maintained high activity despite large differences in temperatures (Female coef*mass*temperature ± SE = −0.121 ± 0.06, z = −2.07, p-value = 0.039, Supplemental Fig. 3, Appendix 4.3).

**Figure 3.**
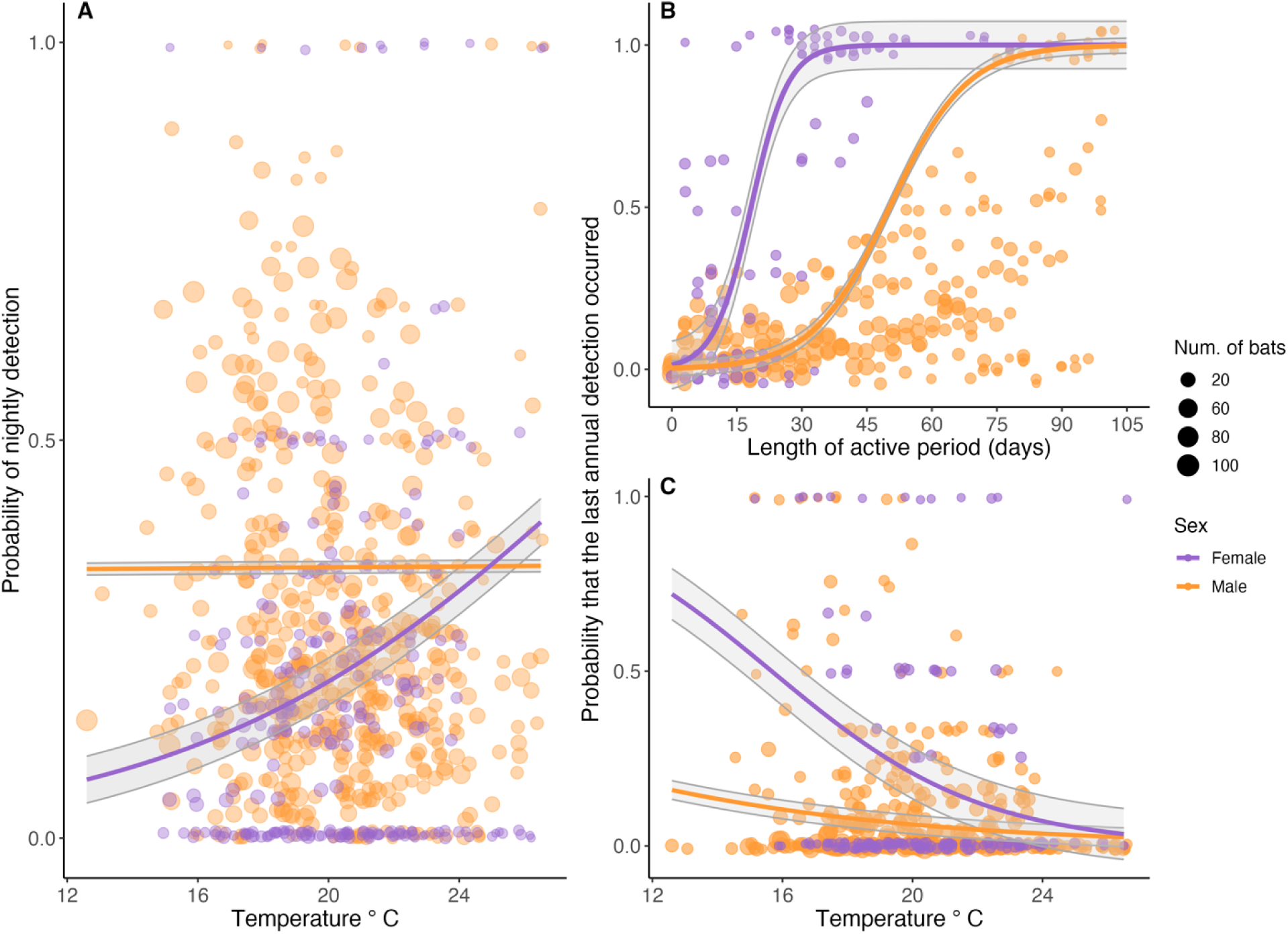
**A.** Effect of temperature on the probability that an individual was active in a night throughout the swarm period. Each point represents the proportion of individuals from a site and year that were active at each temperature by sex. Females are shown in purple and males in orange and lines represent the model predicted probabilities of being active across the temperature gradient. Figure 3B. Relationship between length of the active period in days and the probability of last annual detection for females in purple and males in orange. Lines denote the model predicted probabilities of being detected for the last time within a year. Actual data are binary (1|0) such that 1 = ends activity and 0 = continues to be active. For visualization, points show the proportions of bats of each sex that were detected for the last time within the year, binned across a three-day interval. Figure 3C. Relationship between temperature and probability that bats ended their autumn activity. Each point summarizes the proportion of bats that were detected for the last time at a given temperature for each site and year. Females are shown in purple and males are in orange. Solid lines denote the model predicted probability of being detected for the last time for each sex. Size of points show the number of bats represented in proportion calculations. Notably, males do not end their swarm activity even at temperatures that are relatively low. In each panel, shaded grey regions show standard errors of model predictions.

We also found support for an interaction between the length of the active period and sex on the probability of being detected for the last time (Date intervals since arrival*female = 0.367 ± 0.07, z = 5.49, p-value < 0.001, Fig. 3B; Appendix 2.2), as well as an interactive effect of temperature and sex on the probability of being detected for the last time, such that females were significantly more likely to end their active period as temperatures decreased than males (Temperature*female = −0.164 ± 0.08, z = −2.13, p-value = 0.033, Fig. 3C; Appendix 2.3). Observationally, we also found that within females’ approximate 15-day active period, they were active an average of 4 nights, whereas within males’ 45-day active period they averaged 15 active nights (Supplemental Figure 5, Appendix 4.5).

There was no clear sex bias in pathogen quantity throughout autumn (Female = −0.014 ± 0.10, t = −0.14; p-value = 0.887, Appendix 3.1), however, following autumn swarm, female pathogen loads increased more rapidly, and females had significantly higher pathogen loads than males by early hibernation (Female*date = 0.340 ± 0.08, t = 4.50, p-value < 0.001; Fig. 4A; Appendix 3.2). Ending of autumn activity of males and females generally varied among sites and years and earlier end dates were associated with higher pathogen loads on individuals (Median last date of activity slope on pathogen loads: −1.574 ± 0.41, t = −3.87, p-value < 0.001; Fig. 4B; Appendix 3.3) as well as site-level pathogen quantities in early hibernation (Median last date of activity slope on mean pathogen quantity: −1.562 ± 0.33, t = −4.75, p-value < 0.001; Supplemental Fig. 4; Appendix 4.4). Autumn temperature generally lacked support as an important predictor of pathogen quantity in early hibernation.

**Figure 4.**
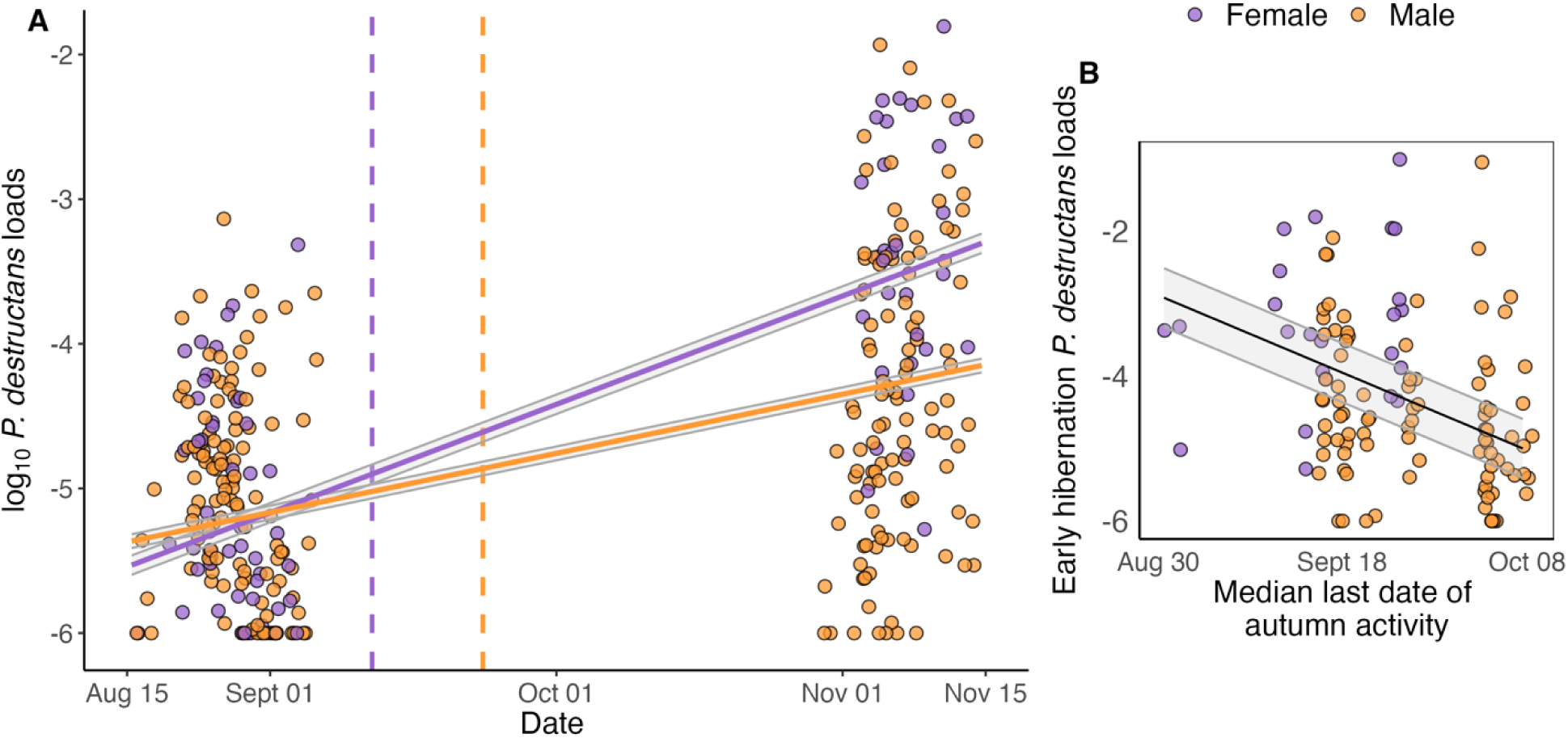
**A:** Pathogen quantity measured on adults of each sex throughout the autumn mating and early hibernation periods. Each point represents an individual bat with females in purple and males in orange. Solid lines denote model predicted fungal loads over time for females and males respectively, with grey shaded regions displaying the standard error. Dashed lines represent the model predicted end dates of activity as estimated in Fig. 2. There was no clear difference in pathogen quantity between sexes during the active mating season in autumn, but female pathogen loads were nearly an order of magnitude higher than male pathogen loads by early hibernation. Figure 4B. Association between the median date of last detection at a site and early hibernation pathogen quantities on individuals. Each point represents an individual bat sampled in early hibernation colored by sex as in panel A and are jittered slightly for visualization. Median last date of autumn activity was calculated for both sexes during each sampling event (i.e., site-year) and used to minimize the outlier effects of a few individuals with exceptionally early or late last active dates. Black line shows model predicted pathogen loads with standard errors displayed with the grey shading. In each panel, date units have been converted to calendar days for clarity.

## 4. Discussion

Sex specific reproductive strategies are common across species, yet how aspects of mating systems give rise to sex-biased disease impacts are often unexplored. Our results suggest that asynchronous swarm phenology between sexes may be a key driver of seasonally sex-biased infection intensity, which are known to affect pathogen transmission and population impacts in other host-pathogen systems (Ferrari et al., 2003; Graham et al., 2013; Grear, Luong, & Hudson, 2012; Perkins, Ferrari, & Hudson, 2008; Sauer et al., 2024; Silk et al., 2018; Skorping & Jensen, 2004). We show female *M. lucifugus* begin swarm activity later and end earlier, spend fewer nights active after arriving to swarm, and have a more pronounced increase in activity on warmer nights compared to males. Importantly, pathogen quantity did not differ between sexes until after the active period of autumn mating, when bats transition to hibernation, and earlier end dates of autumn activity were associated with higher pathogen quantities by early hibernation. Earlier use of torpor by females likely contributes to female-biased infection intensity in early hibernation as it allows an extended period for the pathogen to replicate as opposed to males that remain active and can minimize pathogen replication through euthermia.

In exploring within activity period differences, we found that females are far less active during their shorter swarm period compared to males. This is probably best explained by the need for females to conserve enough energy stores to survive hibernation. Females store sperm overwinter and then give birth and raise offspring following emergence (Czenze, Jonasson, & Willis, 2017; Jonasson & Willis, 2011; Wimsatt, 1945). We provide two new lines of individually derived evidence that aligns with females using energy more conservatively than males. First, we found a clear effect of temperature differently affecting the probability that females and males are active on a given night within their swarm period, and the relationship changed depending on the body condition of females but not males (Supplemental Fig. 3, Appendix 4.3). This suggests that females may be prioritizing the limited days they spend active in autumn to mate and forage on warmer nights that average nearly 20°C (Supplemental Fig. 2B, Appendix 4.2.2) likely because thermoregulating is more efficient at warmer temperatures (Fjelldal et al., 2023; Rubalcaba et al., 2022) and prey resources are more active (Grüebler, Morand, & Naef-Daenzer, 2008; Taylor, 1963). Second, we found that females on average, regardless of their arrival date, begin transitioning into hibernation (i.e., remained in the site/stopped activity) after being active on four nights within their estimated 15 day period, whereas males maintained nightly activity until late in autumn regardless of the date they arrived to swarm, and averaged over 13 active nights (Supplemental Figs 1 and 5, Appendix 4.1 and 4.5) within their 45 day active period. Males that are less constrained by reproductive energy budgets were active more than three times as many nights as females (Supplemental Figs 1 and 5; Appendix 4.1 and 4.5), when temperatures are relatively low (Figs 3A and 3C, Appendix 2.1 and 2.3), and activity was independent of body condition (in contrast to females, Supplemental Fig. 3), suggesting that males are optimizing mating opportunities over energy conservation and supports previous hypotheses (Burns & Broders, 2015; Glover & Altringham, 2008; Rivers, Butlin, & Altringham, 2006; Willis, 2017). Importantly, the disparate reproductive costs between sexes are likely contributing to how each sex interacts with the same novel pressure of an introduced pathogen.

The earlier arrival of males to mating locations than females is observed across many taxa, and two hypotheses have been posed to justify its ubiquity (Morbey & Ydenberg, 2001). First, the ‘rank advantage hypothesis’ states the earliest arriving individuals have access to the best resources or territories, providing a competitive advantage in acquiring females. Second, the ‘mate opportunity hypothesis’ suggests that earlier arriving males can increase the number of females mated, resulting in greater annual reproductive success. In systems under strong sexual selection and in cases where females also have resource-based advantages of arriving early, mate opportunity has more reliably explained the early arrival of males than rank advantage, particularly in species exhibiting sperm competition and highly male-biased sex ratios (Kokko et al., 2006), which is known of temperate bats (Racey et al., 1987; Racey & Entwistle, 2000). Sperm storage by females can also generate advantages for males that more closely synchronize copulations with female receptivity, such that males that mate earlier can secure storage sites in uteri that improve fertilization success (Scialli, 1993). Thus, we hypothesize that the longer swarm period of males is driven by the advantage of increasing copulations with more females in addition to elevating fertilization potential, as suggested by other studies of temperate bats (Burns & Broders, 2015; Pfeiffer & Mayer, 2013). Other environmentally mediated mechanisms likely function in concert with mating advantages. Females migrate from summer maternity colonies and, because foraging to store fat for hibernation is essential for bats in autumn (Kunz, Wrazen, & Burnett, 1998), timing the departure from diminishing resources in summer habitats to optimal resource conditions near swarm locations is likely a more important balance for females that have greater annual energetic constraints than males.

As in other disease systems, sex-specific differences in behavior or physiology during periods of pathogen exposure can result in sex-biased disease impacts and govern host-pathogen dynamics (Guerra-Silveira & Abad-Franch, 2013). We showed that during the season that bats mate and are initially exposed to *P. destructans* after having cleared prior infections (Langwig et al., 2015), activity differs markedly between sexes, however it did not correspond to differences in pathogen quantity within that period (Appendix 3.1) as we expected based on other disease systems (Ferrari et al., 2003; Graham et al., 2013; Guerra-Silveira & Abad-Franch, 2013; Klein, 2000). Furthermore, infections during the autumn mating season or differences in tolerance between sexes are not expected to influence activity patterns because the pathogen quantity on hosts during the active period are likely too low to initiate behaviors or physiological responses symptomatic of disease (Warnecke et al., 2012). Rather, reproductive demands that influence broader phenological patterns are more likely to be shaping sex-biased disease through physiological mechanisms. Female infections rapidly increased almost two orders of magnitude within the two months between the end of their active period and early hibernation (Fig. 4A, Appendix 3.2), suggesting the timing of exposure and pathogen growth likely have temporally decoupled effects on sex-specific disease (Gagnon et al., 2025) in this host-pathogen system. We found that ending activity earlier was associated with higher pathogen loads in early hibernation, which aligns with females being the earliest to hibernate and develop the highest pathogen loads (Fig. 4B, Appendix 3.3). Ending activity is significant in this system because the pathogen can only replicate once bats begin hibernation, and hosts that exceed an infection threshold earlier have lower over-winter survival (Langwig et al., 2016). Consequently, females may be predisposed to have higher disease than males because they initiate hibernation earlier, likely because of their need to conserve energy to achieve reproductive fitness (Czenze et al., 2017; Gagnon et al., 2025).

Phenology evolves to synchronize important biological events, such as reproduction, to optimal conditions and resources (Visser & Gienapp, 2019). However, rapid changes in the biotic or abiotic environment can modify which traits are beneficial or maladaptive (Schlaepfer, Runge, & Sherman, 2002). For bats affected by WNS, if swarm periods evolved to maximize survival and recruitment, disease-induced mortality of early arriving and less active individuals may be selecting for traits that are beneficial for surviving infection but potentially maladaptive for recruitment. Alternatively, if phenology is a plastic trait in bats as it is in other taxa (de Villemereuil et al., 2020), a shift to a later swarm period may minimize disease while still supporting recruitment, thus ameliorating population impacts. Given the clear, primarily additive effect of sex on the timing of activity during the mating period (Fig. 2; Supplemental Table 2; Appendix 4.6.1), we expect that any shifts in activity of one sex will be paralleled by the other. Because females have a more restricted activity period and energy budgets, we expect females to be less flexible in their phenology than males. As such, females will likely need to adapt alternative mechanisms to overcome novel disease pressure, like the reduction in fungal invasion of wing tissue found by Gagnon et al., 2025, or compensatory mechanisms to maintain recruitment if a later swarm window is suboptimal for fitness.

Environmental conditions may also alleviate impacts on females throughout their geographic range or between years. Female bats in milder climates, such as at lower latitudes or years with warmer autumns, may not face the same restriction of suboptimal temperatures that cause bats to begin hibernation. In such instances, females may stay active longer to avoid developing infections severe enough to cause mortality. Similarly, female-biased infections are observed across most host species (Kailing et al., 2023) that share similar reproductive cycles as in Fig. 1. However, more subtle interspecific differences in phenology, like those explored in this study, are largely unknown for most species impacted by WNS but may plausibly contribute to the known variation in early hibernation infections and impacts among species (Langwig et al., 2016; Langwig et al., 2015). Compared to species that mate earlier, species with later swarm activity and delayed hibernation may have lower infections overall that allow more females to survive winter. Previous work has implicated female-biased disease in an initial reduction in the proportion of females following WNS invasion (Kailing et al., 2023), however, many colonies where the pathogen invaded earliest are recovering (Grimaudo et al., 2021; Hoyt, Kilpatrick, & Langwig, 2021; Langwig et al., 2017). Therefore, long-term ecological and evolutionary responses of each sex to disproportionate disease impacts warrant further exploration (Gagnon et al 2025), particularly at spatial and temporal scales beyond this study, as they may contribute to how populations persist.

While sex-based differences in disease have received considerable attention, the mechanisms are often unknown. We demonstrate that sex-specific phenological transitions following exposure can facilitate sex-biased infection intensity, providing a unique example in which physiological asynchronies decouple impacts of the timing of acquiring a pathogen and disease progression between sexes. Importantly, our findings reveal a potentially significant influence of mating dynamics on the trajectory of epidemics and pathways of host persistence or extinction. Other novel pressures that similarly act more strongly on female mating dynamics may impede population recoveries if sex-ratios are disrupted or if traits beneficial for survival are antagonistic for recruitment. Given the importance of demographic structure and recruitment in population impacts, determining how mating strategies influence responses to novel stressors will be critical for conserving species in the face of increasing global change.

## Acknowledgements

We thank Kirsten Fuller and Steffany Yamada for data curation support, and Alexander T. Grimaudo, Nichole A. Laggan, Ariel E. Leon, and Tonie E. Rocke for support in field data collection. We also thank University of Wisconsin Milwaukee Field Station and Gretchen Meyer for site access.

## Author contributions

Macy J. Kailing, Kate E. Langwig, and Joseph R. Hoyt conceived the ideas and designed methodology; Macy J. Kailing, Joseph R. Hoyt, J. Paul White, Jennifer A. Redell, Heather M. Kaarakka, and Kate E. Langwig collected the data; Macy J. Kailing analyzed data with input from Kate E. Langwig and Joseph R. Hoyt. Macy J. Kailing wrote the original manuscript with Kate E. Langwig and Joseph R. Hoyt. All authors contributed critically to the drafts and gave final approval for publication.

## Data availability statement

The datasets generated in this study are available from the Dryad Digital Repository: https://doi.org/10.5061/dryad.rjdfn2zq6 (Kailing et al 2025). Exact site locations are not disclosed to protect endangered species and landowners.

## Supporting information

Additional supporting information may be found in the online version of this article.

Methods, Tables, Figures S1 File detailing supplemental methods, tables, and figures Appendix S1 File containing code and statistical output used throughout the manuscript

