## Supplemental Methods, Tables, and Figures for "Activity patterns during the mating season predict sex-biased infections in an emerging fungal disease"

**File detailing supplemental methods, tables, and figures for:**

**Statistical Methods**

*Directional effect of sex on active period*

To explore whether the direction of the sex effect on swarm periods varied among sampling events, we used Akaike Information Criterion (AIC) to compare models that included site-year as a fixed effect interacting with sex or sex as an additive fixed effect. We compared additive versus interactive sex effects for two model sets; one predicting the start dates of activity and the other predicting end dates of activity (Supplemental Table 2).

*Accounting for sampling bias*

Because many more males were detected than females and we wanted to ensure effects of temperature on nightly activity were not due to this discrepancy, we also tested the GLMM described in Methods 2.4.3 using a balanced dataset truncated by randomly selecting the number of male observations equivalent to the number of female observations from all the male observations (Supplemental Figure 2A). We also used the balanced dataset and explored the relationship between sex and nightly activity using a LMM with temperature as the response. Fixed effects included categorical activity (1 = detected, 0 = undetected) and sex and random effects included site-year and individual (Supplemental Figure 2B).

**Tables**

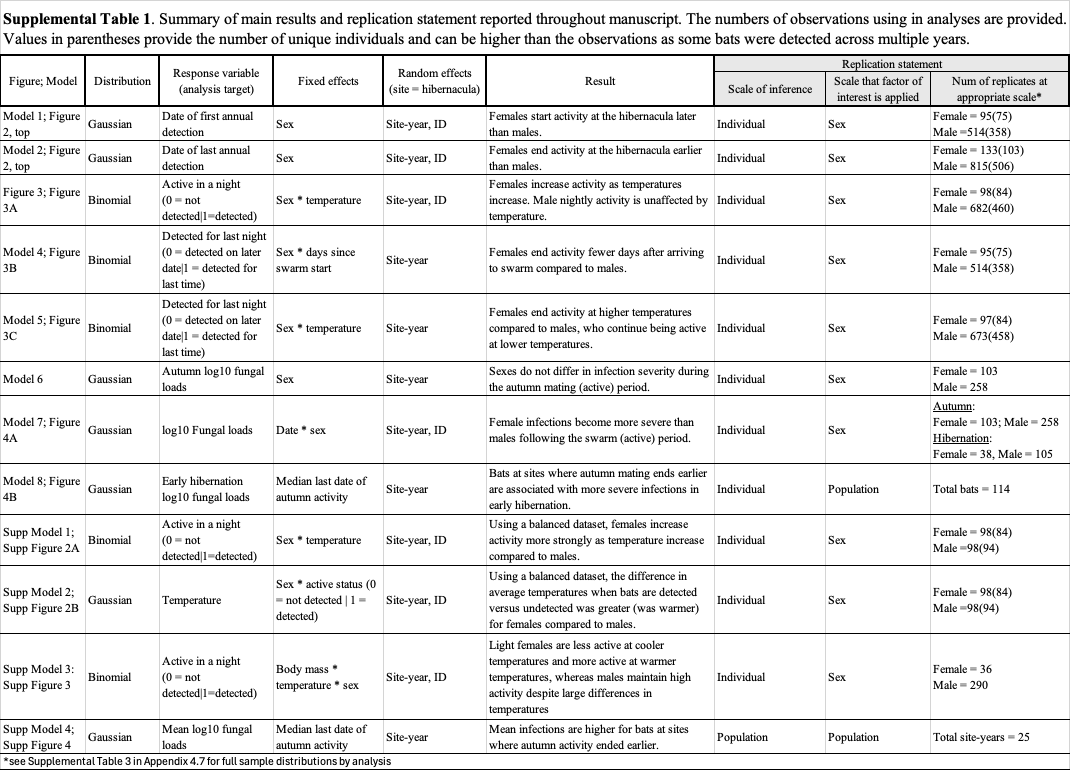

| **Supplemental Table 2:** Results of AIC model comparison testing whether sex is more supported as additive or interactive effect in predicting annual dates of activity across sampling events. Each model includes individual ID as a random effect. | | |
| --- | --- | --- |
| **Response** | **Fixed effects** | **AIC score** |
| First date detected  (e.g. start date) | site-year * sex | 1160.3 |
|  | site-year + sex | 1143.0 |
| Last date detected  (e.g. end date) | site-year * sex | 1711.3 |
|  | site-year + sex | 1693.0 |

**Figures**

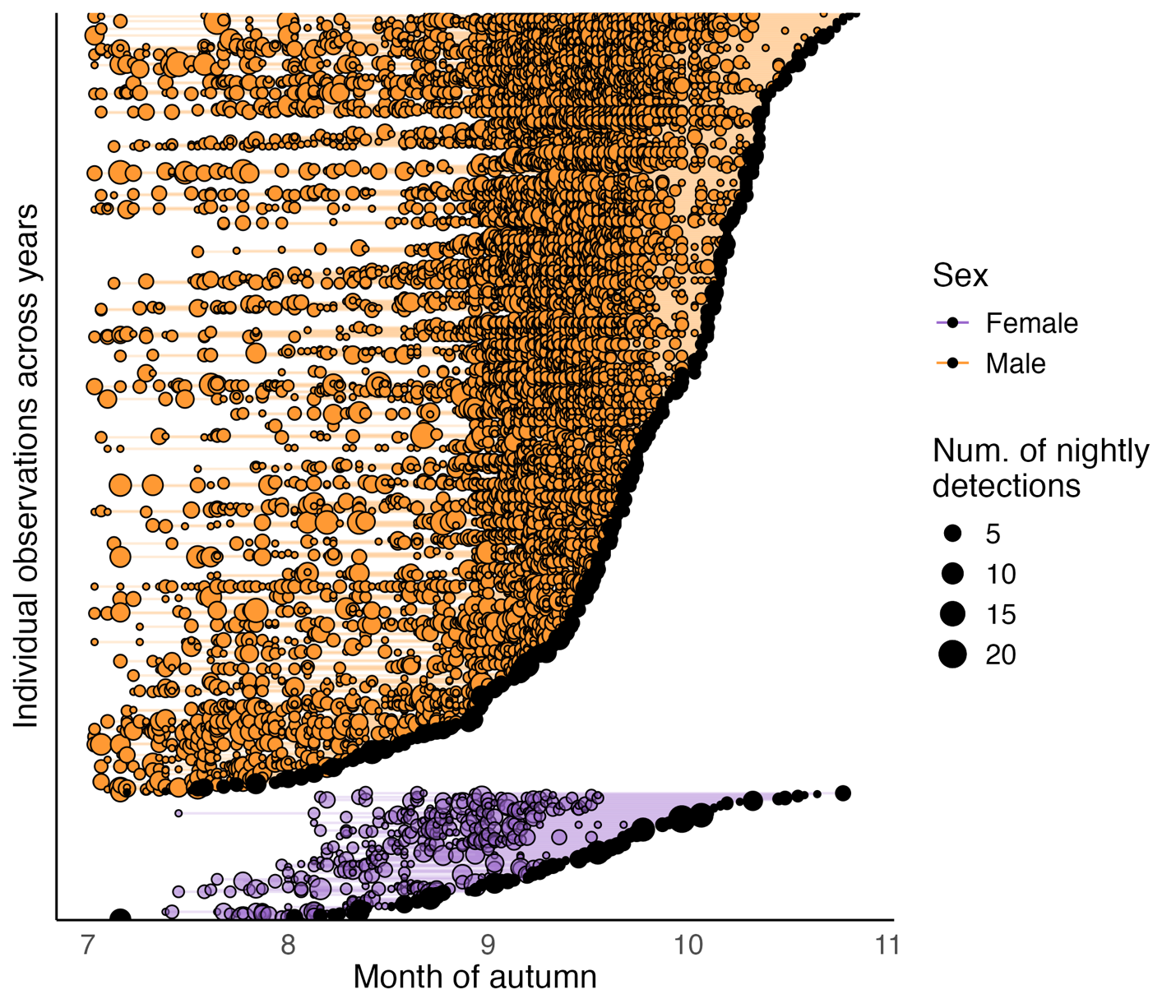

**Supplemental Figure 1:** Overview of individual based activity throughout autumn. Individuals are grouped along the y-axis based on sex then ordered by their last date of activity (black dots). Each point represents a night that an individual was active with the size of points showing the number of times individuals were detected on the respective night.

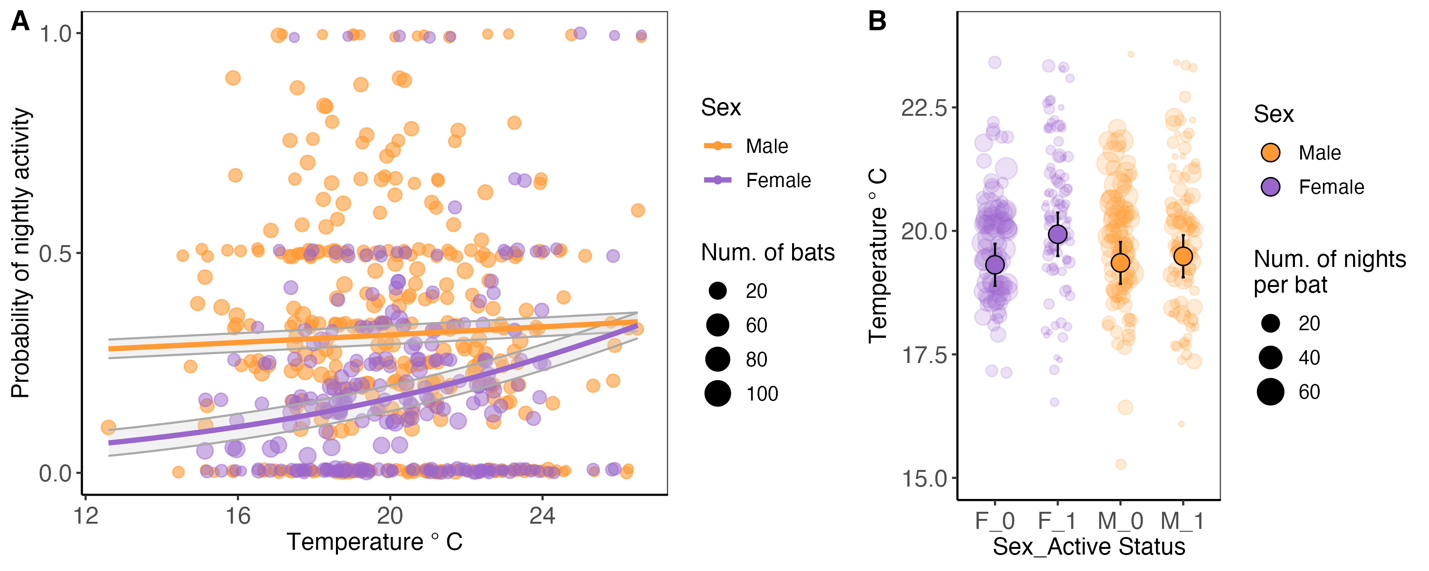

**Supplemental Figure 2A.** Relationship between temperature and the probability of being active by sex after data for males were randomly subsampled to match the number of female observations, ensuring effects were not from sampling bias. Each point represents the proportion of bats that were active at a temperature for each sampling event (site-year) and sized by the number of bats sampled. Females are shown in purple and males in orange. Lines represent model predicted probabilities of being active based on temperature with shaded grey regions displaying standard error. Consistent with Figure 3A, females more strongly increase their activity in response to increasing temperatures. **Supplemental Figure 2B.** Differences in temperature with bat activity. For visualization, points show summarized mean temperatures of individual bats on nights they were active (=1) and undetected (=0) by sex using the balanced dataset. Females and males are shown in purple and orange respectively. Females were present at swarm locations on warmer nights generally, but their active nights occurred on warmer nights relative to their inactive nights. Males had a similar trend of active nights occurring at warmer temperatures relative to nights they were undetected, but to a lesser degree than females.

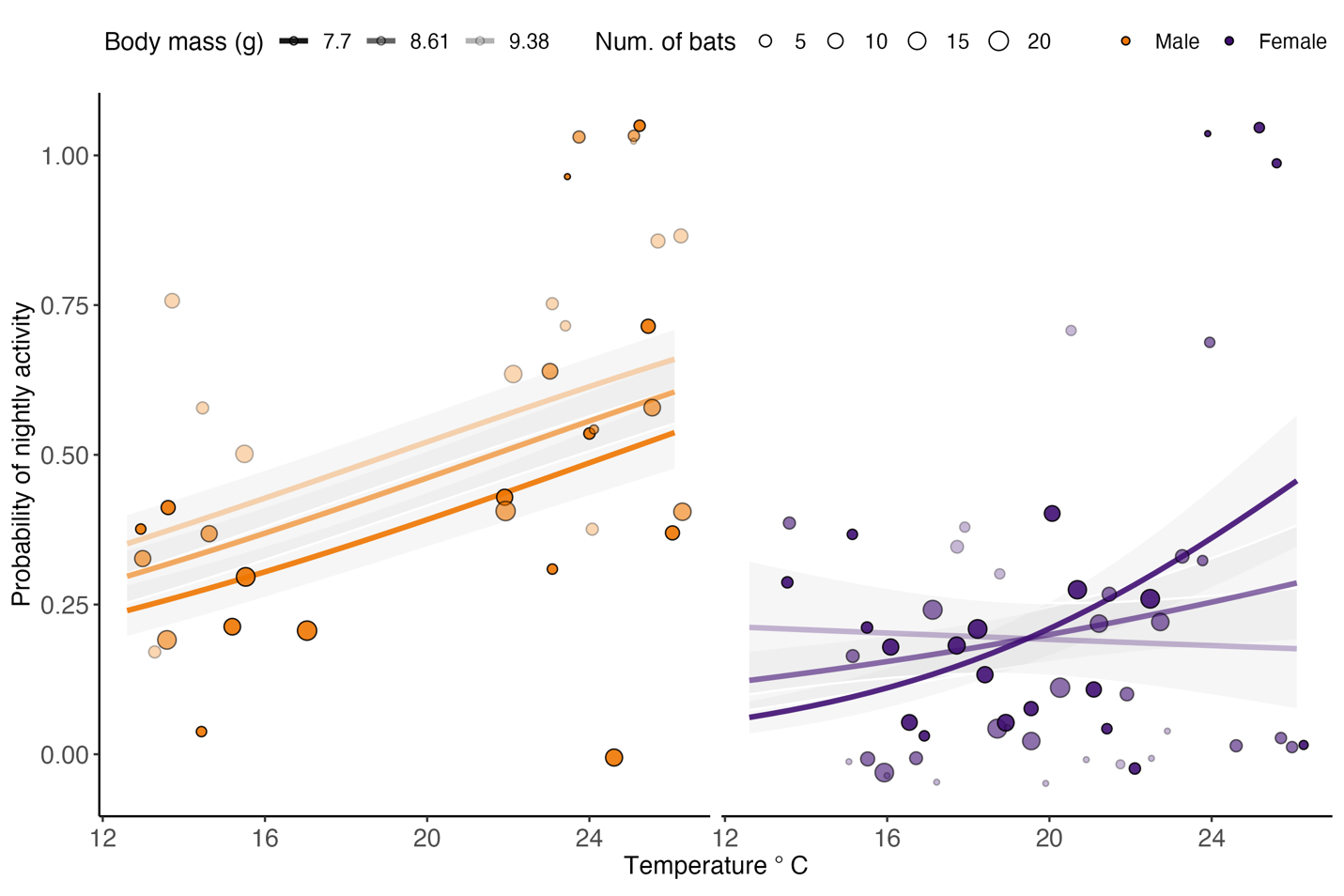

**Supplemental Figure 3.** Effects of body condition and temperature on the probability of nightly activity for each sex. Males are represented by orange on the left and females by purple on the right. Points show the proportions of bats present at the site that were active across the temperature gradient. Proportions were calculated using binned mass and temperature. Lines denote model predicted probability of an individual being active (=1) or undetected (=0) with shaded regions showing the standard error. Body condition affects females differently than males. Lighter females had a higher probability of being active on warmer nights than on cooler nights, indicating that females may adjust activity in accordance with their condition. Male activity was independent of body mass and only slightly positively associated with temperature, suggesting differences in energetic budgeting between sexes.

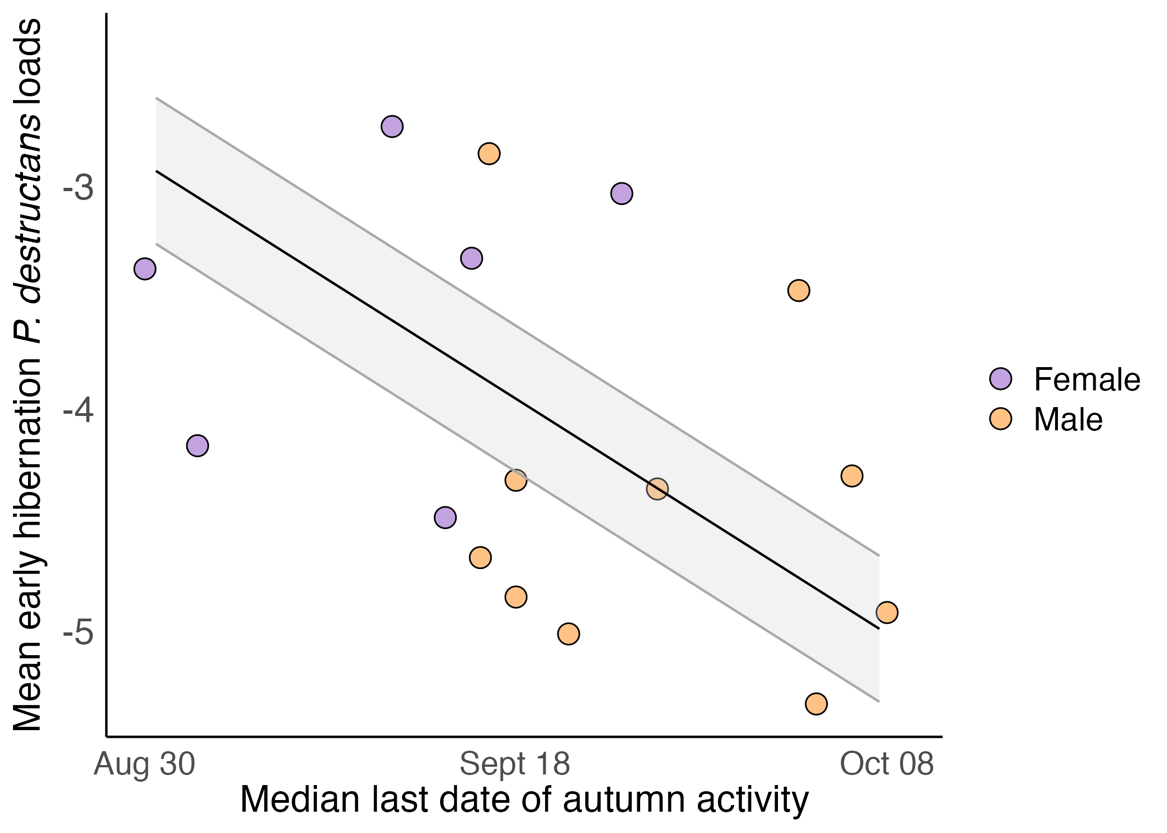

**Supplemental Figure 4**. Relationship between population-level pathogen loads and median ending date of autumn activity. End date of autumn activity was estimated as in Supplemental Figure 4A. Points represent the mean pathogen loads for each sex during the same site-year sampling events that end dates of autumn swarm were characterized. Solid black line denotes model predicted population-level pathogen loads based the ending dates of autumn activity and shaded grey region shows standard error. Fungal loads where swarm activity ceased earlier were generally higher than where bats were active longer.

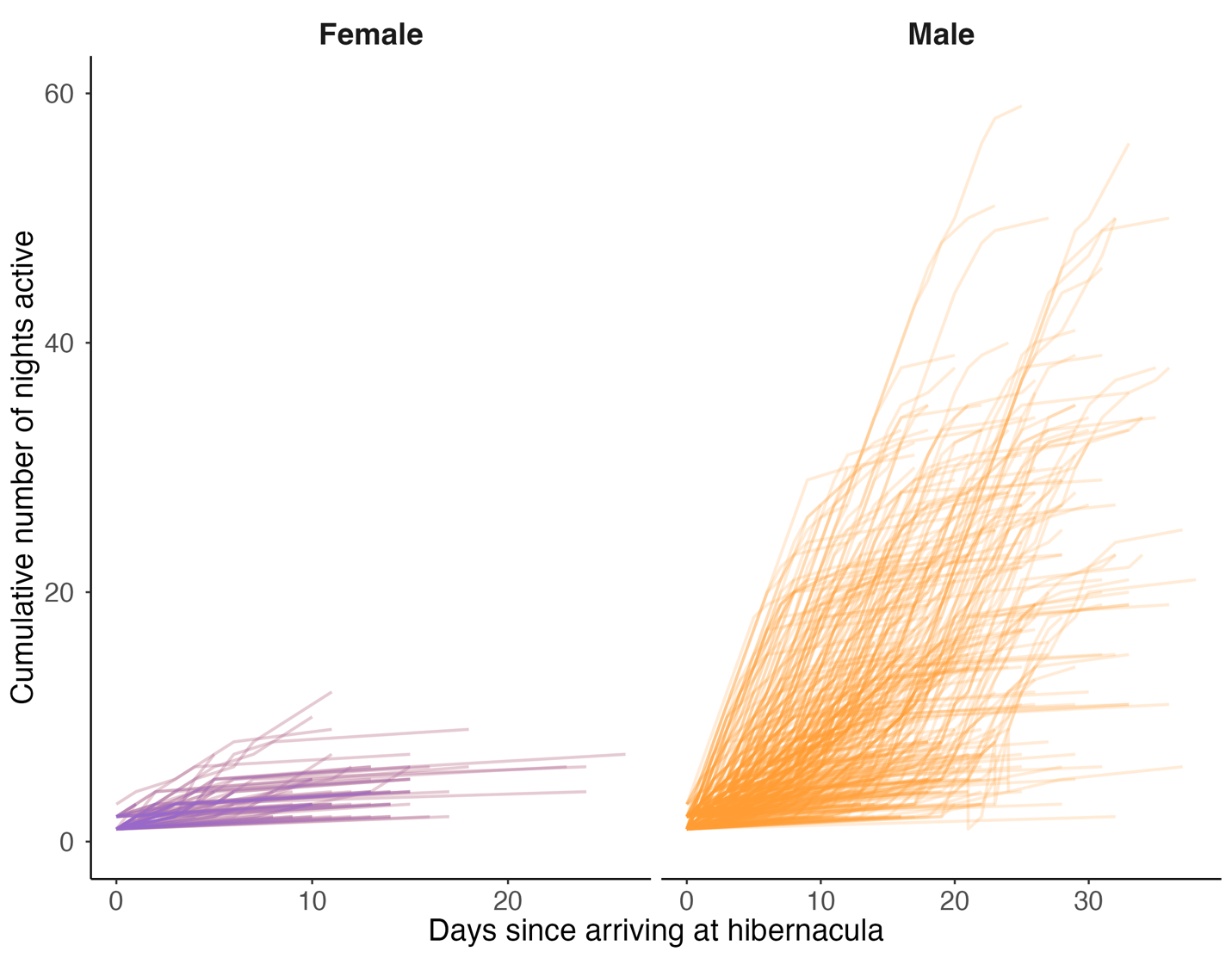

**Supplemental Figure 5**. Individual based cumulative number of nights detected throughout autumn the mating period. Each line represents an individuals’ period of activity (i.e., number of days between the first and last detection). Regardless of variation in when females begin or end swarm, the number of nights they spend active generally plateaus around four. In contrast, males continue to accumulate nights of activity throughout their swarm period, such that no clear maximum threshold of nights active is reached prior to starting hibernation.
