## Supplemental Appendix for "Activity patterns during the mating season predict sex-biased infections in an emerging fungal disease": kailingetal_matingactivity_infection_appendix.nb.html

Appendix for Kailing et al 2025, Mating activity and infection


Code 

- Show All Code
- Hide All Code
- Download Rmd

### Appendix for Kailing et al 2025, Mating activity and infection

##### Activity patterns during the mating season predict sex-biased infections in an emerging fungal disease

###### Macy J. Kailing, Joseph R. Hoyt, J. Paul White, Jennifer A. Redell, Heather M. Kaarakka, and Kate E. Langwig

- 1 When does activity at hibernacula
  during the mating season start and end for each sex? (Figure 2, Top;
  Models 1-2)
  - 1.1 **Model 1**: First,
    test effect of sex on **date of first annual detection**
    (i.e., start of swarm).
  - 1.2 **Model 2**: Then,
    test effect of sex on **date of last annual detection**
    (i.e., end of swarm).
  - 1.3 **Plot Figure 2**.
    Results of active period during autumn mating (Models 1-2)
- 2 What factors contribute to sex
  differences in activity? (Figure 3A-C, Models 3-5)
  - 2.1 **Model 3, Figure
    3A**: First, what are the effects of temperature and sex on
    **nightly activity**?
  - 2.2 **Model 4, Figure
    3B**: Then, what are the effects of the length of the active
    period and sex on the **probability of being detected for the last
    time**?
  - 2.3 **Model 5, Figure
    3C**: Last, what are the effects of temperature and sex on the
    **probability of being detected for the last
    time?**
  - 2.4 **Plot Figure 3**.
    Contribution of temperature and length of swarm on sex-specific activity
    (Models 3-5)
- 3 How does host phenology correspond
  to seasonal infection intensity? (Figure 4A-B, Models 6-8)
  - 3.1 **Model 6**: First,
    does **pathogen quantity** differ between sexes during the
    autumn mating season?
  - 3.2 **Model 7, Figure
    4A**: Then, how does **pathogen quantity** change
    seasonally by sex?
  - 3.3 **Model 8, Figure
    4B**: Last, does the end date of autumn activity affect
    **pathogen quantity in early hibernation**?
  - 3.4 **Plot Figure 4**.
    Seasonal sex-biased pathogen quantity (Figure 4A-B; Models 7-8)
- 4 Supplemental analyses and
  figures
  - 4.1 Supp Figure 1: Visualize complete
    dataset
  - 4.2 Use balanced dataset to support
    sex-specific effects of temperature
    - 4.2.1 Supp Model 1, Supp Fig 2A: Test
      temperature-dependence of bat activity
    - 4.2.2 Supp Model 2, Supp Fig 2B:
      Differences in nightly temperature by sex and active status
    - 4.2.3 Plot Supp Figure 2. Effects of
      temperature with balanced observations between sexes
  - 4.3 Does body condition affect nightly
    activity (Supp Figure 3; Supp Model 3)?
    - 4.3.1 Test whether temperature and
      mass differently influence activity between sexes
    - 4.3.2 Plot Supp Figure 3: Effects of
      mass, temperature and sex on nightly activity
  - 4.4 Additional analysis showing
    association between between phenology and infection intensity (Supp
    Figure 4; Supp Figure 4)
    - 4.4.1 Test whether median end date of
      autumn activity at a site influenced mean pathogen quantity in early
      hibernation
    - 4.4.2 Plot Supp Figure 4. Effect of
      median end date of autumn activity on mean early hibernation pathogen
      quantity at site
  - 4.5 How many total nights are
    individuals active throughout the length of their active period?
    - 4.5.1 Plot Supp Figure 5: Visualize
      cumulative number of active nights by sex
    - 4.5.2 Calculate average number of
      nights detected for each sex
  - 4.6 Model comparisons and
    validation
    - 4.6.1 Compare active period models as
      additive or interactive with site-year and sex as fixed effects (Supp
      Table 2)
    - 4.6.2 View results of null model
      comparisons and AUC estimates from k-fold cross validation of binomial
      models
    - 4.6.3 View AUC scores derived from
      k-fold cross validation of the logistic models reported in the main
      results
  - 4.7 View sampling distributions used
    in each model by site, year and sex where applicable


### 1 When does activity at hibernacula during the mating season start and end for each sex? (Figure 2, Top; Models 1-2)

#### 1.1 **Model 1**: First, test effect of sex on **date of first annual detection** (i.e., start of swarm).


```
m1 = lmer(mindate2~sex + (1|site.wy2) + (1|pit_id),
              control=lmerControl(optimizer="bobyqa", optCtrl=list(maxfun=100000)), 
              data = df.m1); summary(m1)
```


```
Linear mixed model fit by REML ['lmerMod']
Formula: mindate2 ~ sex + (1 | site.wy2) + (1 | pit_id)
   Data: df.m1
Control: lmerControl(optimizer = "bobyqa", optCtrl = list(maxfun = 1e+05))

REML criterion at convergence: 1119.1

Scaled residuals: 
    Min      1Q  Median      3Q     Max 
-3.5190 -0.5320 -0.0346  0.4860  3.9664 

Random effects:
 Groups   Name        Variance Std.Dev.
 pit_id   (Intercept) 0.19965  0.4468  
 site.wy2 (Intercept) 0.01754  0.1324  
 Residual             0.19144  0.4375  
Number of obs: 610, groups:  pit_id, 433; site.wy2, 7

Fixed effects:
            Estimate Std. Error t value
(Intercept)   1.0363     0.0599  17.301
sexF          0.2489     0.0766   3.249

Correlation of Fixed Effects:
     (Intr)
sexF -0.221
```


- ***Result:*** *Females start autumn activity
  at hibernacula later than males.*

#### 1.2 **Model 2**: Then, test effect of sex on **date of last annual detection** (i.e., end of swarm).


```
m2 = lmer(maxdate2~sex + (1|site.wy2) + (1|pit_id),
              control=lmerControl(optimizer="bobyqa", optCtrl=list(maxfun=100000)),
              data = df.m2); summary(m2)
```


```
Linear mixed model fit by REML ['lmerMod']
Formula: maxdate2 ~ sex + (1 | site.wy2) + (1 | pit_id)
   Data: df.m2
Control: lmerControl(optimizer = "bobyqa", optCtrl = list(maxfun = 1e+05))

REML criterion at convergence: 1672.9

Scaled residuals: 
     Min       1Q   Median       3Q      Max 
-2.90593 -0.39386  0.05173  0.51876  2.39627 

Random effects:
 Groups   Name        Variance Std.Dev.
 pit_id   (Intercept) 0.15645  0.3955  
 site.wy2 (Intercept) 0.07805  0.2794  
 Residual             0.20263  0.4501  
Number of obs: 948, groups:  pit_id, 609; site.wy2, 10

Fixed effects:
            Estimate Std. Error t value
(Intercept)  2.71097    0.09313  29.111
sexF        -0.38722    0.06159  -6.287

Correlation of Fixed Effects:
     (Intr)
sexF -0.095
```


- ***Result:*** *Females end autumn activity at
  hibernacula earlier than males.*

#### 1.3 **Plot Figure 2**. Results of active period during autumn mating (Models 1-2)

- Top panel plots data and model coefficients used to determine the
  active period
- Bottom panel visualizes peaks in activity by sex


```
print(p.fig2)
```


- **Key findings:** Females have a shorter active period,
  male activity fully encompasses female activity, and males remain highly
  active later into autumn.

### 2 What factors contribute to sex differences in activity? (Figure 3A-C, Models 3-5)

#### 2.1 **Model 3, Figure 3A**: First, what are the effects of temperature and sex on **nightly activity**?


```
# detect2: 1 = detected | 0 = undetected

m3 = glmer(detect2~bat.tavg*sex + (1|site.wy2) + (1|pit_id), family = binomial(),
            control=glmerControl(optimizer="bobyqa", optCtrl=list(maxfun=100000)),
            data = subset(df.m3)); summary(m3)
```


```
Generalized linear mixed model fit by maximum likelihood (Laplace Approximation) ['glmerMod']
 Family: binomial  ( logit )
Formula: detect2 ~ bat.tavg * sex + (1 | site.wy2) + (1 | pit_id)
   Data: subset(df.m3)
Control: glmerControl(optimizer = "bobyqa", optCtrl = list(maxfun = 1e+05))

     AIC      BIC   logLik deviance df.resid 
 22769.1  22816.7 -11378.5  22757.1    20773 

Scaled residuals: 
    Min      1Q  Median      3Q     Max 
-2.1133 -0.5959 -0.4535  0.7844  4.1081 

Random effects:
 Groups   Name        Variance Std.Dev.
 pit_id   (Intercept) 0.6536   0.8085  
 site.wy2 (Intercept) 0.2089   0.4571  
Number of obs: 20779, groups:  pit_id, 544; site.wy2, 9

Fixed effects:
               Estimate Std. Error z value Pr(>|z|)    
(Intercept)   -0.687705   0.217297  -3.165  0.00155 ** 
bat.tavg       0.001265   0.007507   0.169  0.86618    
sexF          -3.759775   0.627498  -5.992 2.08e-09 ***
bat.tavg:sexF  0.150806   0.030333   4.972 6.64e-07 ***
---
Signif. codes:  0 ‘***’ 0.001 ‘**’ 0.01 ‘*’ 0.05 ‘.’ 0.1 ‘ ’ 1

Correlation of Fixed Effects:
            (Intr) bt.tvg sexF  
bat.tavg    -0.662              
sexF        -0.169  0.233       
bat.tvg:sxF  0.160 -0.241 -0.980
```


```
# view temperature slopes for each sex

kable(m3.emt)
```


| sex | bat.tavg.trend | SE | df | asymp.LCL | asymp.UCL |
| --- | --- | --- | --- | --- | --- |
| M | 0.0012651 | 0.0075070 | Inf | -0.0134483 | 0.0159784 |
| F | 0.1520707 | 0.0294381 | Inf | 0.0943731 | 0.2097683 |


- ***Result:*** *Females are less active
  generally throughout the mating period, but female activity increases
  with temperature. Male activity is less affected by
  temperature.*

#### 2.2 **Model 4, Figure 3B**: Then, what are the effects of the length of the active period and sex on the **probability of being detected for the last time**?


```
# last.det2: 1 = last day active | 0 = detected on subsequent nights
# isa: date intervals since arrival

m4 = glmer(last.det2~isa*sex + (1|site.wy2) + (1|pit_id), family=binomial(),
           control=glmerControl(optimizer="bobyqa", optCtrl=list(maxfun=100000)),
           data=subset(df.m4));summary(m4)
```


```
Generalized linear mixed model fit by maximum likelihood (Laplace Approximation) ['glmerMod']
 Family: binomial  ( logit )
Formula: last.det2 ~ isa * sex + (1 | site.wy2) + (1 | pit_id)
   Data: subset(df.m4)
Control: glmerControl(optimizer = "bobyqa", optCtrl = list(maxfun = 1e+05))

     AIC      BIC   logLik deviance df.resid 
  2800.5   2839.4  -1394.3   2788.5     4787 

Scaled residuals: 
   Min     1Q Median     3Q    Max 
-1.665 -0.300 -0.142 -0.043 74.209 

Random effects:
 Groups   Name        Variance Std.Dev.
 pit_id   (Intercept) 5.469    2.339   
 site.wy2 (Intercept) 1.602    1.266   
Number of obs: 4793, groups:  pit_id, 433; site.wy2, 7

Fixed effects:
            Estimate Std. Error z value Pr(>|z|)    
(Intercept) -4.29912    0.70400  -6.107 1.02e-09 ***
isa          0.70560    0.07336   9.618  < 2e-16 ***
sexM        -1.37247    0.52423  -2.618  0.00884 ** 
isa:sexM    -0.36702    0.06682  -5.492 3.97e-08 ***
---
Signif. codes:  0 ‘***’ 0.001 ‘**’ 0.01 ‘*’ 0.05 ‘.’ 0.1 ‘ ’ 1

Correlation of Fixed Effects:
         (Intr) isa    sexM  
isa      -0.535              
sexM     -0.603  0.501       
isa:sexM  0.507 -0.950 -0.622
```


- ***Result:*** *Regardless of arrival date,
  the length of female active periods are shorter than males.*

#### 2.3 **Model 5, Figure 3C**: Last, what are the effects of temperature and sex on the **probability of being detected for the last time?**


```
# last.det2: 1 = last day active | 0 = detected on subsequent nights

m5 = glmer(last.det2~bat.tavg*sex + (1|pit_id) + (1|site.wy2), family = binomial(),
            control=glmerControl(optimizer="bobyqa", optCtrl=list(maxfun=100000)),
            data = subset(df.m5)); summary(m5)
```


```
Generalized linear mixed model fit by maximum likelihood (Laplace Approximation) ['glmerMod']
 Family: binomial  ( logit )
Formula: last.det2 ~ bat.tavg * sex + (1 | pit_id) + (1 | site.wy2)
   Data: subset(df.m5)
Control: glmerControl(optimizer = "bobyqa", optCtrl = list(maxfun = 1e+05))

     AIC      BIC   logLik deviance df.resid 
  2647.9   2687.9  -1318.0   2635.9     5780 

Scaled residuals: 
    Min      1Q  Median      3Q     Max 
-1.1904 -0.2993 -0.2391 -0.1375  9.1936 

Random effects:
 Groups   Name        Variance Std.Dev.
 pit_id   (Intercept) 0.01401  0.1184  
 site.wy2 (Intercept) 0.51982  0.7210  
Number of obs: 5786, groups:  pit_id, 542; site.wy2, 9

Fixed effects:
              Estimate Std. Error z value Pr(>|z|)    
(Intercept)    0.16202    0.56715   0.286  0.77512    
bat.tavg      -0.14453    0.02669  -5.416 6.11e-08 ***
sexF           4.67270    1.54263   3.029  0.00245 ** 
bat.tavg:sexF -0.16407    0.07701  -2.131  0.03313 *  
---
Signif. codes:  0 ‘***’ 0.001 ‘**’ 0.01 ‘*’ 0.05 ‘.’ 0.1 ‘ ’ 1

Correlation of Fixed Effects:
            (Intr) bt.tvg sexF  
bat.tavg    -0.895              
sexF        -0.284  0.314       
bat.tvg:sxF  0.286 -0.321 -0.994
```


- ***Result:*** *Females end activity on warmer
  nights compared to males that remain active even as temperature
  decreases.*

#### 2.4 **Plot Figure 3**. Contribution of temperature and length of swarm on sex-specific activity (Models 3-5)


```
print(p.fig3)
```


- **Key findings**: Female activity is generally reduced
  compared to males and the probability of nightly activity increases with
  temperature. Females also end activity fewer days after arriving at
  hibernacula and at higher temperatures.

### 3 How does host phenology correspond to seasonal infection intensity? (Figure 4A-B, Models 6-8)

#### 3.1 **Model 6**: First, does **pathogen quantity** differ between sexes during the autumn mating season?


```
m6 = lmer(lgdL2~sex + (1|site.wy2), 
          data = df.m6); summary(m6)
```


```
Linear mixed model fit by REML ['lmerMod']
Formula: lgdL2 ~ sex + (1 | site.wy2)
   Data: df.m6

REML criterion at convergence: 314.2

Scaled residuals: 
    Min      1Q  Median      3Q     Max 
-1.8127 -0.5967 -0.1576  0.5203  3.5268 

Random effects:
 Groups   Name        Variance Std.Dev.
 site.wy2 (Intercept) 0.2147   0.4634  
 Residual             0.3193   0.5650  
Number of obs: 172, groups:  site.wy2, 9

Fixed effects:
            Estimate Std. Error t value
(Intercept) -5.35210    0.17072 -31.350
sexF        -0.01397    0.09788  -0.143

Correlation of Fixed Effects:
     (Intr)
sexF -0.159
```


- ***Result:*** *No clear difference in
  pathogen loads between sexes during the autumn mating season.*

#### 3.2 **Model 7, Figure 4A**: Then, how does **pathogen quantity** change seasonally by sex?


```
m7 = lmer(lgdL2~sex*pdate2 + (1|site.wy2), 
          data = df2); summary(m7)
```


```
Linear mixed model fit by REML ['lmerMod']
Formula: lgdL2 ~ sex * pdate2 + (1 | site.wy2)
   Data: df2

REML criterion at convergence: 822

Scaled residuals: 
    Min      1Q  Median      3Q     Max 
-2.0626 -0.6530 -0.0952  0.6261  3.5533 

Random effects:
 Groups   Name        Variance Std.Dev.
 site.wy2 (Intercept) 0.2592   0.5091  
 Residual             0.5722   0.7564  
Number of obs: 343, groups:  site.wy2, 13

Fixed effects:
             Estimate Std. Error t value
(Intercept) -11.90983    0.69417 -17.157
sexM          3.06440    0.76088   4.027
pdate2        0.74914    0.06673  11.227
sexM:pdate2  -0.34035    0.07560  -4.502

Correlation of Fixed Effects:
            (Intr) sexM   pdate2
sexM        -0.754              
pdate2      -0.971  0.776       
sexM:pdate2  0.748 -0.992 -0.783
```


- ***Result:*** *Females develop higher
  pathogen loads than males by early hibernation.*

#### 3.3 **Model 8, Figure 4B**: Last, does the end date of autumn activity affect **pathogen quantity in early hibernation**?


```
m8 = lmer(lgdL2~med.maxdate + (1|site.wy2), 
           control=lmerControl(optimizer="bobyqa", optCtrl=list(maxfun=100000)),
           data = df.m8); summary(m8)
```


```
Linear mixed model fit by REML ['lmerMod']
Formula: lgdL2 ~ med.maxdate + (1 | site.wy2)
   Data: df.m8
Control: lmerControl(optimizer = "bobyqa", optCtrl = list(maxfun = 1e+05))

REML criterion at convergence: 299.9

Scaled residuals: 
    Min      1Q  Median      3Q     Max 
-2.1009 -0.5816 -0.0953  0.6783  3.2574 

Random effects:
 Groups   Name        Variance Std.Dev.
 site.wy2 (Intercept) 0.6249   0.7905  
 Residual             0.6642   0.8150  
Number of obs: 114, groups:  site.wy2, 10

Fixed effects:
            Estimate Std. Error t value
(Intercept)  11.1252     3.9794   2.796
med.maxdate  -1.5740     0.4071  -3.867

Correlation of Fixed Effects:
            (Intr)
med.maxdate -0.998
```

#### 3.4 **Plot Figure 4**. Seasonal sex-biased pathogen quantity (Figure 4A-B; Models 7-8)


```
print(p.fig4)
```


- **Key findings:**
  - Female-biased infection intensity only arises following the active
    autumn mating period.
  - Infection intensity is higher on bats associated with sites where
    activity ends earlier.

### 4 Supplemental analyses and figures

#### 4.1 Supp Figure 1: Visualize complete dataset


```
print(p.sf1)
```


- Overall, males are more active than females throughout autumn and
  fully encompass female swarm activity.

#### 4.2 Use balanced dataset to support sex-specific effects of temperature

##### 4.2.1 Supp Model 1, Supp Fig 2A: Test temperature-dependence of bat activity


```
sm1 = glmer(detect2~bat.tavg*sex + (1|site.wy2) + (1|pit_id), family = binomial(),
             control=glmerControl(optimizer="bobyqa", optCtrl=list(maxfun=100000)),
             data = subset(df1.tru)); summary(sm1)
```


```
Generalized linear mixed model fit by maximum likelihood (Laplace Approximation) ['glmerMod']
 Family: binomial  ( logit )
Formula: detect2 ~ bat.tavg * sex + (1 | site.wy2) + (1 | pit_id)
   Data: subset(df1.tru)
Control: glmerControl(optimizer = "bobyqa", optCtrl = list(maxfun = 1e+05))

     AIC      BIC   logLik deviance df.resid 
  4354.4   4392.6  -2171.2   4342.4     4266 

Scaled residuals: 
    Min      1Q  Median      3Q     Max 
-1.5418 -0.5403 -0.4081 -0.2618  4.1241 

Random effects:
 Groups   Name        Variance Std.Dev.
 pit_id   (Intercept) 0.5480   0.7402  
 site.wy2 (Intercept) 0.0166   0.1288  
Number of obs: 4272, groups:  pit_id, 178; site.wy2, 8

Fixed effects:
              Estimate Std. Error z value Pr(>|z|)    
(Intercept)   -1.19533    0.43121  -2.772  0.00557 ** 
bat.tavg       0.02066    0.02126   0.972  0.33107    
sexF          -3.17607    0.74400  -4.269 1.96e-05 ***
bat.tavg:sexF  0.11847    0.03614   3.279  0.00104 ** 
---
Signif. codes:  0 ‘***’ 0.001 ‘**’ 0.01 ‘*’ 0.05 ‘.’ 0.1 ‘ ’ 1

Correlation of Fixed Effects:
            (Intr) bt.tvg sexF  
bat.tavg    -0.964              
sexF        -0.573  0.565       
bat.tvg:sxF  0.563 -0.582 -0.980
```


```
#view temperature slopes for each sex

kable(sm1.emt)
```


| sex | bat.tavg.trend | SE | df | asymp.LCL | asymp.UCL |
| --- | --- | --- | --- | --- | --- |
| M | 0.0206591 | 0.0212553 | Inf | -0.0210005 | 0.0623187 |
| F | 0.1391326 | 0.0293756 | Inf | 0.0815574 | 0.1967078 |

##### 4.2.2 Supp Model 2, Supp Fig 2B: Differences in nightly temperature by sex and active status


```
# detect2 (active status): 1=detected | 0=undetected

sm2 = lmer(bat.tavg~detect2*sex + (1|pit_id) + (1|site.wy2), 
           data = subset(df1.tru)); summary(sm2)
```


```
Linear mixed model fit by REML ['lmerMod']
Formula: bat.tavg ~ detect2 * sex + (1 | pit_id) + (1 | site.wy2)
   Data: subset(df1.tru)

REML criterion at convergence: 19136

Scaled residuals: 
    Min      1Q  Median      3Q     Max 
-3.1646 -0.6908 -0.0348  0.7159  3.1546 

Random effects:
 Groups   Name        Variance Std.Dev.
 pit_id   (Intercept) 0.3333   0.5773  
 site.wy2 (Intercept) 1.3436   1.1591  
 Residual             4.9435   2.2234  
Number of obs: 4272, groups:  pit_id, 178; site.wy2, 8

Fixed effects:
              Estimate Std. Error t value
(Intercept)   19.35260    0.42333  45.715
detect21       0.13314    0.10616   1.254
sexF          -0.03922    0.12987  -0.302
detect21:sexF  0.48511    0.17724   2.737

Correlation of Fixed Effects:
            (Intr) dtct21 sexF  
detect21    -0.086              
sexF        -0.138  0.264       
detct21:sxF  0.051 -0.600 -0.322
```

##### 4.2.3 Plot Supp Figure 2. Effects of temperature with balanced observations between sexes


```
print(p.sf2)
```


- ***Using the balanced dataset between sexes, results are
  consistent with Model 3 in Figure 3A:***
  - *Female nightly activity increases with temperature, but male
    activity is relatively unaffected by temperature.*
  - *The difference between mean temperature on nights when bats are
    active compared to nights when bats are undetected is greater for
    females than males.*
  - *Females concentrate their activity to warmer nights, whereas
    males do not.*

#### 4.3 Does body condition affect nightly activity (Supp Figure 3; Supp Model 3)?

##### 4.3.1 Test whether temperature and mass differently influence activity between sexes


```
sm3 <- glmmTMB(detect2~sex*mass*temp + (1|pit_id) + (1|site.wy2),
               family = binomial(),
               data = subset(df.sm3)); summary(sm3)
```


```
 Family: binomial  ( logit )
Formula:          detect2 ~ sex * mass * temp + (1 | pit_id) + (1 | site.wy2)
Data: subset(df.sm3)

     AIC      BIC   logLik deviance df.resid 
  5113.5   5177.1  -2546.8   5093.5     4248 

Random effects:

Conditional model:
 Groups   Name        Variance Std.Dev.
 pit_id   (Intercept) 0.6763   0.8223  
 site.wy2 (Intercept) 0.2774   0.5267  
Number of obs: 4258, groups:  pit_id, 336; site.wy2, 10

Conditional model:
                 Estimate Std. Error z value Pr(>|z|)  
(Intercept)     -4.972624   2.307187  -2.155   0.0311 *
sexF           -18.493191   9.659752  -1.915   0.0556 .
mass             0.338061   0.257427   1.313   0.1891  
temp             0.106098   0.117137   0.906   0.3651  
sexF:mass        2.046204   1.164068   1.758   0.0788 .
sexF:temp        1.026604   0.485330   2.115   0.0344 *
mass:temp       -0.001248   0.013107  -0.095   0.9242  
sexF:mass:temp  -0.121309   0.058665  -2.068   0.0387 *
---
Signif. codes:  0 ‘***’ 0.001 ‘**’ 0.01 ‘*’ 0.05 ‘.’ 0.1 ‘ ’ 1
```

##### 4.3.2 Plot Supp Figure 3: Effects of mass, temperature and sex on nightly activity


```
print(p.sf3)
```


- ***Results:***
  - *Females in lower body condition (ie body mass) concentrate
    activity to warm nights, but females in higher body condition do
    not.*
  - *Males were more active generally, but body condition did not
    affect the temperatures at which they were active.*
  - *Sexes are likely budgeting energy differently.*

#### 4.4 Additional analysis showing association between between phenology and infection intensity (Supp Figure 4; Supp Figure 4)

##### 4.4.1 Test whether median end date of autumn activity at a site influenced mean pathogen quantity in early hibernation


```
sm4 = lmer(mean.lgdL~med.maxdate + (1|site.wy2), data = subset(df2.sum)); summary(sm4)
```


```
Linear mixed model fit by REML ['lmerMod']
Formula: mean.lgdL ~ med.maxdate + (1 | site.wy2)
   Data: subset(df2.sum)

REML criterion at convergence: 31.1

Scaled residuals: 
     Min       1Q   Median       3Q      Max 
-0.82223 -0.53563 -0.06433  0.47551  1.18360 

Random effects:
 Groups   Name        Variance Std.Dev.
 site.wy2 (Intercept) 0.5788   0.7608  
 Residual             0.1053   0.3244  
Number of obs: 16, groups:  site.wy2, 10

Fixed effects:
            Estimate Std. Error t value
(Intercept)  11.0041     3.1930   3.446
med.maxdate  -1.5615     0.3285  -4.753

Correlation of Fixed Effects:
            (Intr)
med.maxdate -0.997
```

##### 4.4.2 Plot Supp Figure 4. Effect of median end date of autumn activity on mean early hibernation pathogen quantity at site


```
print(p.sf4)
```


- ***Result:*** *Consistent with Figure 4B,
  infection intensity in early hibernation is greater at sites where
  autumn activity ends earlier.*

#### 4.5 How many total nights are individuals active throughout the length of their active period?

##### 4.5.1 Plot Supp Figure 5: Visualize cumulative number of active nights by sex


```
print(p.sf5)
```

##### 4.5.2 Calculate average number of nights detected for each sex


```
kable(n.dates.sum)
```


| sex | mean\_n.dates |
| --- | --- |
| F | 3.810526 |
| M | 13.947471 |


- Males are active for ~14 nights on average and continue activity
  without a clear threshold, whereas females cease activity after an
  average of ~4 nights.

#### 4.6 Model comparisons and validation

##### 4.6.1 Compare active period models as additive or interactive with site-year and sex as fixed effects (Supp Table 2)

First, begin date of activity models


```
m1b.add = lmer(mindate2~site.wy2 + sex + (1|pit_id), data = df.m1)
m1b.int = lmer(mindate2~site.wy2 * sex + (1|pit_id), data = df.m1)
```


- Compare using AIC


```
kable(AIC(m1b.add,m1b.int))
```


|  | df | AIC |
| --- | --- | --- |
| m1b.add | 10 | 1142.960 |
| m1b.int | 16 | 1160.287 |


Then, end date of activity models


```
m2b.add = lmer(maxdate2~site.wy2 + sex + (1|pit_id), data = df.m2)
m2b.int = lmer(maxdate2~site.wy2 * sex + (1|pit_id), data = df.m2)
```


- Compre using AIC


```
kable(AIC(m2b.add,m2b.int))
```


|  | df | AIC |
| --- | --- | --- |
| m2b.add | 13 | 1692.986 |
| m2b.int | 20 | 1711.309 |


- ***Results***
  - *Sex is more supported as an additive rather than interactive
    effect in models including site-year as fixed effects.*
  - *The dates that activity starts and ends varies among sites and
    years, but females are consistently later to start and earlier to end
    compared to males.*

##### 4.6.2 View results of null model comparisons and AUC estimates from k-fold cross validation of binomial models


```
kable(nullcomps_full)
```


|  | df | AIC |
| --- | --- | --- |
| m1.null | 4 | 1134.2857 |
| m1 | 5 | 1129.1469 |
| m2.null | 4 | 1715.0409 |
| m2 | 5 | 1682.9043 |
| m3.null | 3 | 22821.9059 |
| m3 | 6 | 22769.0835 |
| m4.null | 2 | 3623.1434 |
| m4 | 6 | 2800.5278 |
| m5.null | 2 | 2741.8440 |
| m5 | 6 | 2647.9294 |
| m7.null | 3 | 966.0277 |
| m7 | 6 | 833.9688 |
| m8.null | 3 | 319.9266 |
| m8 | 4 | 307.8518 |


- All AICs of reported models are >2 scores below the null models
  (structured with response~1), indicating improvement over the null.

##### 4.6.3 View AUC scores derived from k-fold cross validation of the logistic models reported in the main results


```
kable(kfold_full)
```


| Model | AUC |
| --- | --- |
| m3 | 69.11% |
| m4 | 78.35% |
| m5 | 84.82% |


- Results indicate that 70%, 78%, and 85% of our test data was
  successfully predicted by the training models of Model 3, Model 4, and
  Model 5, respectively.

#### 4.7 View sampling distributions used in each model by site, year and sex where applicable


|  |  |  |
| --- | --- | --- |
| **ST3A. Model 1 Sample summary: Start dates of activity** | | |
| Year | Females | Males |
| BA IT | | |
| --- | --- | --- |
| 2021 | 11 | 60 |
| 2022 | 16 | 134 |
| MA N | | |
| 2021 | 13 | 33 |
| 2022 | 10 | 72 |
| NE MI | | |
| 2021 | 2 | 36 |
| 2022 | 17 | 88 |
| 2023 | 26 | 92 |

|  |  |  |
| --- | --- | --- |
| **ST3B. Model 2 Sample summary: End dates of activity** | | |
| Year | Females | Males |
| BA IT | | |
| --- | --- | --- |
| 2020 | 4 | 63 |
| 2021 | 12 | 102 |
| 2022 | 17 | 139 |
| MA N | | |
| 2020 |  | 10 |
| 2021 | 18 | 73 |
| 2022 | 16 | 98 |
| NE MI | | |
| 2020 |  | 20 |
| 2021 | 7 | 73 |
| 2022 | 24 | 107 |
| 2023 | 35 | 130 |

|  |  |  |
| --- | --- | --- |
| **ST3C. Model 3 Sample summary: Prob of nightly activity** | | |
| Year | Females | Males |
| BA IT | | |
| --- | --- | --- |
| 2020 | 4 | 63 |
| 2021 | 12 | 101 |
| 2022 | 17 | 139 |
| MA N | | |
| 2020 |  | 10 |
| 2021 | 18 | 71 |
| 2022 | 16 | 100 |
| NE MI | | |
| 2020 |  | 20 |
| 2021 | 7 | 73 |
| 2022 | 24 | 107 |

|  |  |  |
| --- | --- | --- |
| **ST3D. Model 4 Sample summary: Prob of last detection** | | |
| Year | Females | Males |
| BA IT | | |
| --- | --- | --- |
| 2021 | 11 | 60 |
| 2022 | 16 | 134 |
| MA N | | |
| 2021 | 13 | 33 |
| 2022 | 10 | 73 |
| NE MI | | |
| 2021 | 2 | 36 |
| 2022 | 17 | 88 |
| 2023 | 26 | 92 |

|  |  |  |
| --- | --- | --- |
| **ST3E. Model 5 Sample summary: Prob of last detection** | | |
| Year | Females | Males |
| BA IT | | |
| --- | --- | --- |
| 2020 | 4 | 63 |
| 2021 | 12 | 97 |
| 2022 | 17 | 139 |
| MA N | | |
| 2020 |  | 10 |
| 2021 | 18 | 67 |
| 2022 | 16 | 100 |
| NE MI | | |
| 2020 |  | 20 |
| 2021 | 6 | 72 |
| 2022 | 24 | 107 |

|  |  |  |  |
| --- | --- | --- | --- |
| **ST3F. Model 7 Sample summary: Seasonal infections** | | | |
| Year | Season | Females | Males |
| BA IT | | | |
| --- | --- | --- | --- |
| 2019 | Early hiber | 10 | 9 |
| 2020 | Autumn mating | 26 | 66 |
| 2020 | Early hiber | 5 | 12 |
| 2021 | Autumn mating |  | 3 |
| 2021 | Early hiber | 1 | 10 |
| 2022 | Autumn mating | 11 | 38 |
| 2022 | Early hiber | 3 | 13 |
| MA N | | | |
| 2019 | Early hiber | 4 | 15 |
| 2020 | Autumn mating | 12 | 28 |
| 2020 | Early hiber | 5 | 11 |
| 2021 | Autumn mating | 14 | 12 |
| 2021 | Early hiber | 2 | 7 |
| 2022 | Autumn mating | 22 | 48 |
| 2022 | Early hiber | 4 | 9 |
| NE MI | | | |
| 2019 | Early hiber | 5 | 13 |
| 2020 | Autumn mating | 5 | 37 |
| 2020 | Early hiber | 1 | 11 |
| 2021 | Autumn mating |  | 6 |
| 2021 | Early hiber | 5 | 8 |
| 2022 | Autumn mating | 23 | 34 |
| 2022 | Early hiber | 10 | 6 |
| 2023 | Early hiber |  | 20 |

|  |  |
| --- | --- |
| **ST3G. Model 8 Sample summary: Phenology-dependent early hibernation infections** | |
| Year | N |
| BA IT | |
| --- | --- |
| 2020 | 16 |
| 2021 | 8 |
| 2022 | 10 |
| MA N | |
| 2020 | 8 |
| 2021 | 9 |
| 2022 | 10 |
| NE MI | |
| 2020 | 7 |
| 2021 | 12 |
| 2022 | 15 |
| 2023 | 19 |
